# Gene editing without *ex vivo* culture evades genotoxicity in human hematopoietic stem cells

**DOI:** 10.1101/2023.05.27.542323

**Authors:** Jing Zeng, My Anh Nguyen, Pengpeng Liu, Lucas Ferreira da Silva, Linda Y. Lin, David G. Justus, Karl Petri, Kendell Clement, Shaina N. Porter, Archana Verma, Nola R. Neri, Tolulope Rosanwo, Marioara-Felicia Ciuculescu, Daniela Abriss, Esther Mintzer, Stacy A. Maitland, Selami Demirci, John F. Tisdale, David A. Williams, Lihua Julie Zhu, Shondra M. Pruett-Miller, Luca Pinello, J. Keith Joung, Vikram Pattanayak, John P. Manis, Myriam Armant, Danilo Pellin, Christian Brendel, Scot A. Wolfe, Daniel E. Bauer

## Abstract

Gene editing the *BCL11A* erythroid enhancer is a validated approach to fetal hemoglobin (HbF) induction for β-hemoglobinopathy therapy, though heterogeneity in edit allele distribution and HbF response may impact its safety and efficacy. Here we compared combined CRISPR-Cas9 endonuclease editing of the *BCL11A* +58 and +55 enhancers with leading gene modification approaches under clinical investigation. We found that combined targeting of the *BCL11A* +58 and +55 enhancers with 3xNLS-SpCas9 and two sgRNAs resulted in superior HbF induction, including in engrafting erythroid cells from sickle cell disease (SCD) patient xenografts, attributable to simultaneous disruption of core half E-box/GATA motifs at both enhancers. We corroborated prior observations that double strand breaks (DSBs) could produce unintended on-target outcomes in hematopoietic stem and progenitor cells (HSPCs) such as long deletions and centromere-distal chromosome fragment loss. We show these unintended outcomes are a byproduct of cellular proliferation stimulated by ex vivo culture. Editing HSPCs without cytokine culture bypassed long deletion and micronuclei formation while preserving efficient on-target editing and engraftment function. These results indicate that nuclease editing of quiescent hematopoietic stem cells (HSCs) limits DSB genotoxicity while maintaining therapeutic potency and encourages efforts for in vivo delivery of nucleases to HSCs.

## INTRODUCTION

The β-hemoglobinopathies, including SCD and transfusion-dependent β-thalassemia (TDT), are the most prevalent inherited monogenic disorders worldwide. Globally, there are an estimated 300,000 births with SCD and 25,000 with TDT every year^1, 2^. SCD is an autosomal recessive severe multisystem disorder caused by a single A-to-T base substitution in the β-globin gene *HBB*, resulting in a missense mutation of glutamic acid to valine at amino acid 6. The resulting hemoglobin S (α2β^S^2) polymerizes when deoxygenated, damages the erythrocyte, and leads to hemolytic anemia, vaso-occlusion, inflammation, and hypercoagulability with recurrent pain and progressive multi-organ injury. In TDT, insufficient production of β-globin, caused by more than 400 mutations in *HBB*, results in an imbalanced α/β-chain ratio. The precipitation of excessive free α-globin chains impairs the maturation of erythroid precursors and produces ineffective erythropoiesis, chronic hemolysis, and compensatory hematopoietic expansion. Hydroxyurea, supportive care, regular blood transfusions, and iron chelation are current mainstay treatments for SCD and TDT. Curative allogeneic bone marrow transplantation is limited by availability of suitable donors and immune incompatibility. Autologous hematopoietic stem cell transplantation is emerging as gene therapy and gene editing methods advance^3, 4^.

Fetal hemoglobin (HbF, α2γ2) is the best established modifier of β-hemoglobinopathy clinical severity^5^. Fetuses and newborns are protected by high HbF levels with disease only beginning to manifest in infancy as levels wane. Individuals with common and rare genetic variants associated with elevated HbF have milder disease. Currently, there are several gene therapy and gene editing clinical trials for SCD and TDT intended to reactivate HbF based on shRNA knockdown of the BCL11A repressor in erythroid precursors and CRISPR-Cas nuclease or base editor mediated disruption of the *BCL11A* erythroid specific +58 enhancer or BCL11A binding sequences at the *HBG1/HBG2* γ-globin promoters (NCT03745287, NCT03655678, NCT04211480, NCT04853576, NCT05456880). Although initial trials have shown encouraging results with HbF induction, lessening of anemia and improvement of clinical features, long-term outcomes in large patient cohorts have yet to be ascertained^6, 7^. HbF levels vary widely at baseline among individuals, suggesting potential heterogeneity in HbF responses after gene therapy, analogous to heterogeneity in HbF responses observed after pharmacotherapy^8^.

Several reports have indicated that individuals with compound heterozygous sickle hemoglobin (HbS) and hereditary persistence of fetal hemoglobin (HPFH), with pancellularly distributed HbF levels in the ∼30-45% range, are much more mildly affected than typical SCD but can still infrequently present with SCD complications such as vaso-occlusive episodes, acute chest syndrome, osteonecrosis, hemiparesis, retinal changes, and need for intermittent blood transfusions^9–14^. HbF ∼30-45% is a similar range of HbF level as has been observed in the trials of autologous gene therapy^6, 7^. Prior calculations using a mathematical model of Hb polymerization and *in vitro* experiments have suggested that more HbF, ∼50% or higher, may be required to achieve inhibition of deoxyHbS polymerization similar to that calculated or observed in *in vitro* studies for RBCs from patients with heterozygous sickle cell trait HbAS, where the fraction of HbA is ∼65% *in vivo*^15, 16^.

Beyond the β-hemoglobin disorders, gene editing has tremendous promise to remedy a wide range of monogenic and complex diseases^17^. Gene editing typically relies on a programmable endonuclease to produce a DSB followed by endogenous cellular repair. After a genetic modification is introduced, further rounds of cleavage cease and the genetic change (“edit”) becomes fixed in the genome. There are a wide diversity of possible endogenous DNA damage repair pathways^18^. The major DNA repair pathway is non-homologous end-joining (NHEJ), which is active in all phases of the cell cycle. Minor pathways that may be harnessed for desired genome editing outcomes include microhomology-mediated end joining^19^ (MMEJ) and, in the presence of an extrachromosomal donor DNA template sequence, homology directed repair (HDR)^20^. Additional potential DSB repair outcomes include long deletions with extensive unidirectional end resection^21^ and centromere distal chromosome arm loss which may result in micronuclei and chromothripsis^22^. Clinical trials of HSC editing to date have not reported the comprehensive distribution of allelic outcomes after gene editing. Here we explore the hypothesis that a modified procedure could both increase HbF induction potency and decrease genotoxic potential of gene editing.

## RESULTS

### Combined *BCL11A* +58 and +55 enhancer editing maximizes HbF induction

Prior studies have revealed two strong erythroid enhancers of *BCL11A* contained in the +58 and +55 DNase I hypersensitive sites (DHSs), with GATA1 binding, erythroid chromatin accessibility, heterologous enhancer potential, and sensitivity to loss-of-function^23–27^. Indels at a core GATA1 binding site at the *BCL11A* +58 enhancer lead to increased HbF levels in erythrocytes in preclinical models and in clinical trials^6, 24, 28–30^. We used SpCas9:sgRNA ribonucleoprotein (RNP) electroporation to compare the impact of gene editing at the +58 and +55 enhancers individually or in combination. We observed that +58 enhancer edited erythroid precursors (edited with sgRNA #1617, aka sg1617) lose chromatin accessibility at the +58 enhancer but accessibility remains at the +55 enhancer, suggesting residual enhancer function preserving partial *BCL11A* expression (**Figure 1A**). To investigate functional sequences at the *BCL11A* +55 erythroid enhancers, we electroporated CD34+ HSPCs with 3xNLS-SpCas9 protein and 15 modified synthetic chimeric single guide RNAs (sgRNAs) targeting positions within core sequences at the +55 enhancer (**Figure S1A**), as defined by prior functional screening and chromatin profiling^24, 25, 31, 32^. Each of the RNPs led to highly efficient editing, with mean indels ranging from 94.9%-99.9% (**Figure S1B**). Following erythroid differentiation culture, hemoglobin HPLC showed that the greatest HbF induction was associated with editing by sgRNAs sg1449 and sg1450, which produced indels disrupting a half E-box/GATA (TGN7-9WGATAR) motif, the binding site for GATA1 and TAL1 erythroid transcriptional activation complexes^33, 34^ (**Figures S1A, S1C, S1D** and **S1E**). Of note, 3xNLS-SpCas9 editing with sg1617 targeting the *BCL11A* +58 enhancer, which leads to potent HbF induction including in therapeutic gene editing clinical trials^6, 30^ also generates indels that disrupt a half E-box/GATA motif^24, 28, 29^. Gene editing with sgRNA #1450, aka sg1450, produced partial reduction of +55 chromatin accessibility, but preserved +58 chromatin accessibility in primary erythroid precursors (**Figure 1A**).

**Figure 1.**
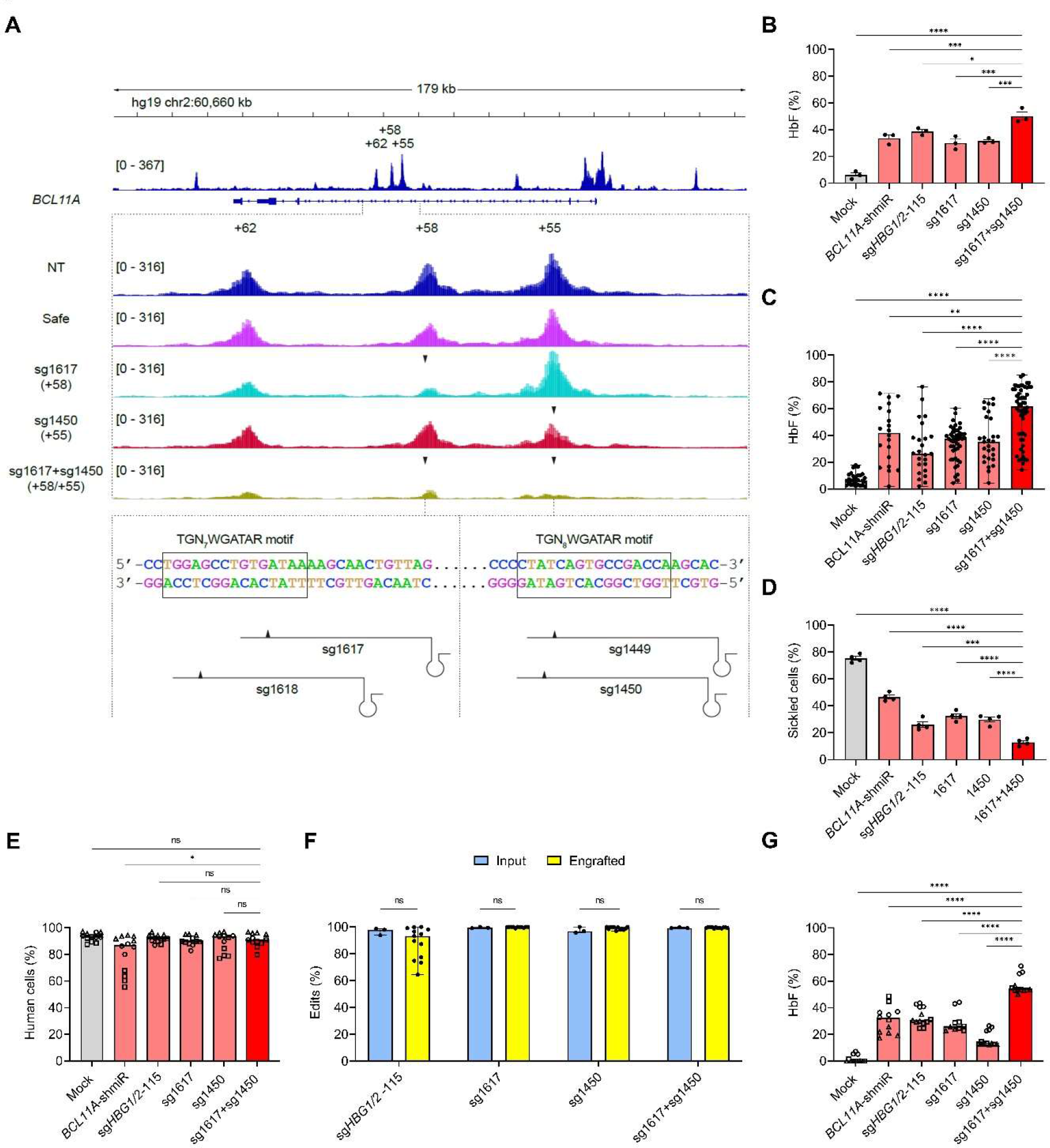
Robust HbF induction by combined editing of the +58 and +55 *BCL11A* erythroid enhancers in human CD34+ HSPCs (A) ATAC-seq analysis of chromatin accessibility following single editing of *BCL11A* +58 or +55 *BCL11A* enhancer, and combined editing of +58 and +55 *BCL11A* enhancers in human CD34+ HSPC derived primary erythroid precursors from three healthy donors (HD 1, HD 2 and HD 3). Schema shown sgRNAs targeting +58 and +55 *BCL11A* enhancer with the cleavage site relative to the TGN_7-9_WGATAR (B) HbF induction by HPLC analysis in CD34+ HSPC derived *in vitro* differentiated erythroid cells from one healthy donor and two SCD patients (HD 7, SCD 1 and SCD 2). Data are plotted as mean ± SEM and analyzed with one-way analysis of variance (ANOVA). \**P* < 0.5, \*\*\**P* < 0.001, \*\*\*\**P* < 0.0001. (C) HbF level by HPLC from individual erythroid single cell derived clonal liquid cultures derived from single CD34+ HSPCs from one healthy donor and two SCD patients (HD 8, SCD 1 and SCD 2). Data are plotted as median with range and analyzed with one-way ANOVA. \*\**P* < 0.01, \*\*\*\**P* < 0.0001. (D) Fraction of sickled cells from SCD *in vitro* differentiated enucleated erythroid cells by MBS sickling assay. Data were from two SCD donors (SCD 1 and SCD 2) and two technical replicates for each donor. Data are plotted as mean ± SEM and analyzed with one-way ANOVA. \*\*\**P* < 0.001, \*\*\*\**P* < 0.0001. (E) Human chimerism 16 weeks post transplantation in bone marrow. Data are plotted as grand median and analyzed with one-way ANOVA. ns: nonsignificant, \**P* < 0.5. n = 11-13 primary recipients. Donors were from HD 7, SCD 1 and SCD 2. (F) Indel frequencies were measured by Sanger sequencing for sg*HBG*1/2-115, sg1617 and sg1450. Overall editing was measured by +58 drop-off ddPCR assay for sg1617+sg1450. Data are plotted as median with range and analyzed with the unpaired two-tailed Student’s *t*-test. ns: nonsignificant. n = 3 donors (HD 7, SCD 1 and SCD 2) for input, n = 11-13 primary recipients for engrafted. (G) HbF levels by HPLC analysis in engrafted erythroid cells. Data are plotted as grand median and analyzed with one-way ANOVA. \*\*\*\**P* < 0.0001. n = 11-13 primary recipients. Donors were from HD 7, SCD 1 and SCD 2.

We hypothesized that combined editing targeting both the +58 and +55 enhancers could lead to greater enhancer disruption, loss of erythroid *BCL11A* expression, and HbF induction as compared to editing a single enhancer alone. Indeed we found that chromatin accessibility was greatly reduced at both +55 and +58 enhancers, as well as at the adjacent +62 enhancer, after combined editing with 3xNLS-SpCas9 and both sg1617 and sg1450 sgRNAs (**Figure 1A**).

Targeting the +58, +55, or both enhancers resulted in 56.4%, 63.9%, or 75.5% reduction in *BCL11A* mRNA level (**Figure S1F**). We compared *in vitro* erythroid maturation in control cells, those edited at the +58 enhancer alone (with sg1617), the +55 enhancer alone (with sg1450), or at both the +58 and +55 enhancers (sg1617+sg1450). We found similar cell expansion and erythroid maturation *in vitro*, including enucleation frequency with each editing approach (**Figures S1G and S1H**). Consistent with more potent inhibition of *BCL11A* expression, targeting both enhancers resulted in substantially greater HbF induction as compared to targeting either enhancer alone, with 70.2%, 76.8%, and 91.7% F-cells and 39.7%, 43.4%, 62.2% HbF tetramer by HPLC (**Figures S1I and S1J**). We also evaluated two other clinically relevant approaches for HbF induction, microRNA embedded shRNA targeting the *BCL11A* transcript selectively in erythroid precursors (aka *BCL11A*-shmiR)^7, 35^, and introducing edits to the *HBG1* and *HBG2* promoters targeting binding sequences around position -115 that recruit BCL11A and are the site of naturally occurring HPFH mutations (aka *HBG1/2* -115 editing)^36–39^.

We found that combined targeting of the *BCL11A* +58 and +55 enhancers with 3xNLS-SpCas9 and two sgRNAs resulted in the most potent HbF induction (50.1%±5.4%) of tested approaches as compared to *BCL11A* +58 editing alone (30.1%±4.9%), *BCL11A* +55 editing alone (31.6±1.9%), *HBG1/2* promoter editing (38.7%±2.6%), *BCL11A*-shmiR (33.7%±4.2%), or mock (6.0%±2.9%) (**Figure 1B**). We isolated single cell derived erythroid colonies from each of these conditions and found that combined editing of the +58 and +55 enhancers produced greater median HbF level (61.9%) as compared to single +58 or +55 enhancer editing (37.5% or 35.4% respectively), *HBG1/2* -115 editing (26.5%), or BCL11A-shmiR (41.9%) (**Figure 1C**). From SCD donors, we sorted enucleated erythroid cells and exposed the cells to sodium metabisulfite to evaluate deoxygenation-induced hemoglobin polymerization-associated shape change (i.e. sickled cells)^28^. We observed fewer sickled cells following combined +58 and +55 enhancer editing as compared to any of the other HbF induction approaches (**Figure 1D** and **Figure S2A**).

We compared combined +58 and +55 *BCL11A* enhancer editing to individual +58 or +55 *BCL11A* editing *in vivo*, by infusing RNP edited CD34+ HSPCs to immunodeficient NBSGW mice, which permit human multilineage engraftment without conditioning therapy^40^. We observed similar human chimerism in the recipients of +58 single edited, +55 single edited, *HBG1/2* -115 edited HSPCs, and modestly reduced chimerism in *BCL11A*-shmiR transduced HSPCs (**Figure 1E**). Since we infused equal numbers of cells from each condition, but *BCL11A*-shmiR transduced cells showed more proliferation during *ex vivo* culture (0.7-fold, 0.78-fold, 0.64-fold, 0.84-fold, and 1.4-fold for combined +58/+55 editing, single +58 editing, single +55 editing, *HBG1/2* -115 editing, and *BCL11A*-shmiR transduction respectively), a reduced fraction of infused HSCs might account for this reduced chimerism. Gene modification was robust across conditions, with median 99.5% overall gene edits for combined +58/+55 editing, 100% gene edits for single +58 editing, 99.3**%** for single +55 editing, 93.2% for *HBG1/2* -115 editing, and vector copy number 3.2 for *BCL11A*-shmiR transduced engrafting cells (**Figures 1F** and **S2B**).

We observed the greatest HbF induction for combined +58/+55 gene editing (with 53.8% in edited engrafting BM erythroid cells) as compared to single +58 editing (26.3%), single +55 editing (13.2%), *HBG1/2* -115 editing (31.2%), *BCL11A*-shmiR transduced engrafting cells (35.3%), and unedited cells (0.28%) (**Figure 1G**). We observed similar multilineage engraftment of B-lymphoid cells, granulocytes, monocytes, and HSPCs across all the conditions (**Figures S2C** and **S2D**). *BCL11A*-shmiR, single +58 enhancer editing and combined +58/+55 *BCL11A* enhancer editing had lower erythroid repopulation as compared to mock (**Figure S2E**), although no difference was observed in the fraction of enucleated erythroid cells in HD and SCD recipients (**Figure S2F**). *BCL11A* expression in sorted BM erythroid cells was reduced in the gene edited compared to the control group (**Figure S2G**). *BCL11A* expression in B cells, HSPCs and non-B, non-HSPC, non-erythroid cells was similar between mock and combined +58/+55 enhancers edited groups (**Figures S2H**, **S2I** and **S2J**). We infused bone marrow cells from primary recipients into secondary recipients and evaluated bone marrow after an additional 16 weeks. We observed combined enhancer edited and unedited HSPCs maintained similar secondary repopulation potential consistent with editing of long-term HSCs (**Figure S2K**).

### Combined enhancer editing disrupts two critical motifs

We predicted that producing simultaneous DSBs at the +58 and +55 enhancers would produce programmed 3.1 kb deletions and 3.1 kb inversions in addition to indels at each cleavage site^41–43^. We developed ddPCR assays to specifically detect each of these expected edits. In addition, we utilized a ddPCR drop-off assay where any edit, including 1 bp insertions or deletions, results in loss of signal. To distinguish short indels from longer deletions or rearrangements, we designed a ddPCR offset assay where only unedited alleles and short indel alleles preserve signal (**Figures 2A** and **S2L**-**S2O**). We found that a median of 67.8% of alleles consisted of programmed 3.1 kb deletions and inversions in input cells. A modest increase in programmed deletions and inversions (median 83.2%) and corresponding decrease in short indels was observed in engrafting cells as compared to input (**Figure S2P**). The observed decrease in short indels in engrafting cells was due to a decline in MMEJ-mediated indels with no change in the fraction of NHEJ-mediated indels (**Figure S2Q**). A high frequency of edits was preserved in secondary engrafting cells (**Figure S2R**).

**Figure 2.**
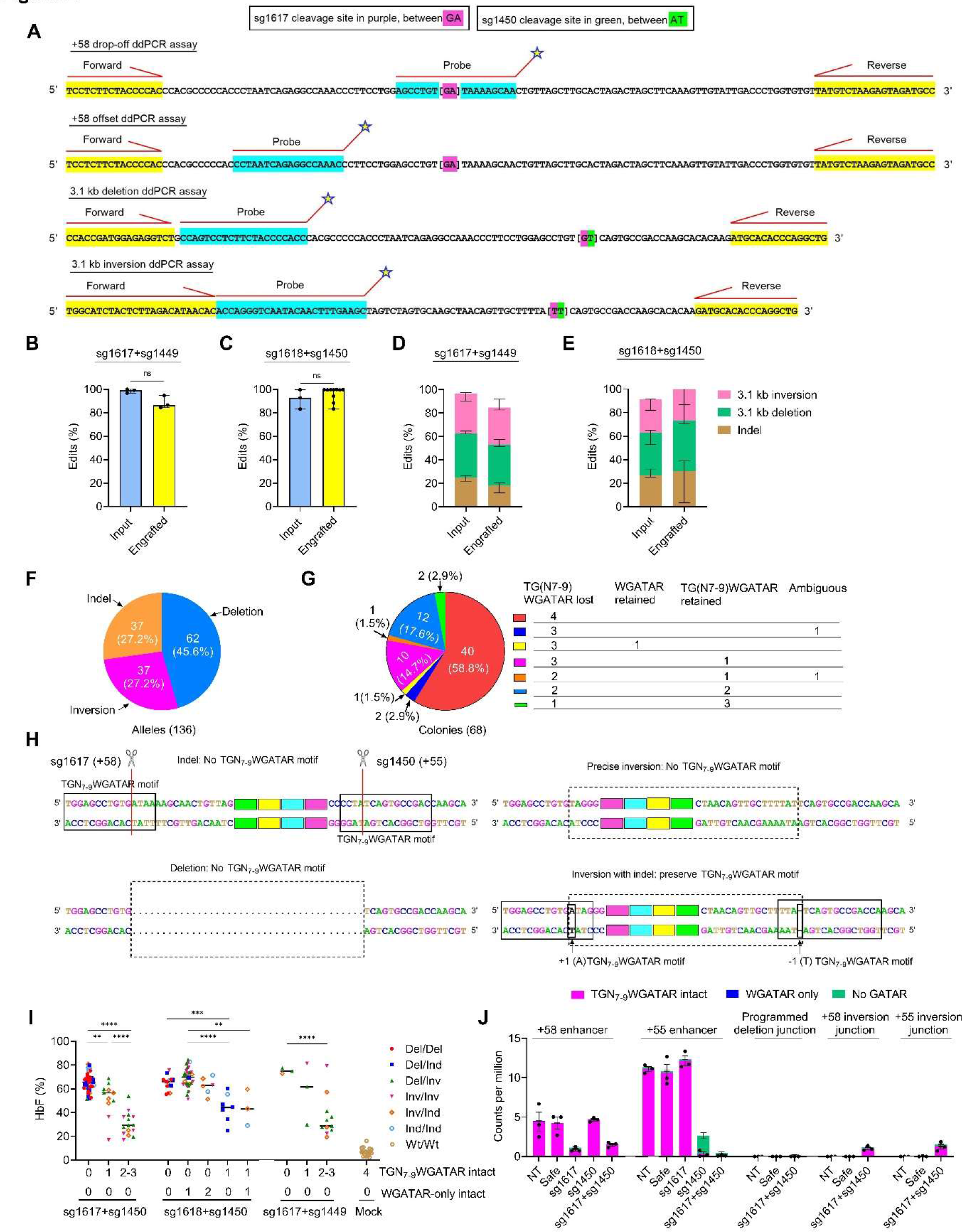
Potent disruption of TGN7-9WGATAR half E-box/GATA binding motifs after combined editing of the +58 and +55 *BCL11A* erythroid enhancers in human HSPCs (A) Design of primers and probe for ddPCR assays to quantify overall edit by +58 drop-off ddPCR assay, small indel by +58 offset ddPCR assay, 3.1 kb deletion and 3.1 kb inversion by 3.1 kb deletion ddPCR assay and 3.1 kb inversion ddPCR assay, respectively. (B and C) Overall edit in engrafted cells compared to input cells for sg1617+sg1449 editing (B) by +58 sg1617 drop-off ddPCR assay and sg1618+sg1450 editing (C) by +58 sg1618 drop-off ddPCR assay. Data are plotted as median with range and analyzed with the unpaired two-tailed Student’s *t*-test. ns: nonsignificant. n = 3 healthy donors (HD 16, HD 17 and HD 18 for sg1617+sg1449, HD 7, HD13 and HD 14 for sg1617+sg1450) for input, n = 3-10 primary recipients for engrafted. (D and E) Frequencies of indels, deletion and inversion of sg1617+sg1449 (D) and sg1618+sg1450 (E) by deletion, inversion, +58 offset and +58 drop-off ddPCR assays in engrafted cells compared to input cells. Data are plotted as median with range. n = 3 healthy donors (HD 16, HD 17 and HD 18 for sg1617+sg1449, HD 7, HD13 and HD 14 for sg1617+sg1450) for input, n = 3-10 primary recipients for engrafted. (F) Allelic analysis of single clones after sg1617+sg1450 editing of CD34+ HSPCs from HD 8. (G) Junctional analysis of TGN7-9WGATAR half E-box/GATA binding motifs after sg1617+sg1450 editing in individual single cell derived erythroid colonies after editing of CD34+ HSPCs from HD 8. See schematic panel H for example editing outcomes. (H) Schema of edit outcomes after sg1617+sg1450 editing and its associated disruption/preservation of TGN7-9WGATAR half E-box/GATA binding motifs. (I) Genotypes and junction analysis of TGN7-9WGATAR half E-box/GATA binding motifs compared to HbF levels by HPLC from colonies subject to combined editing of the +58 and +55 *BCL11A* erythroid enhancers with three distinct sgRNA pairs. Data are plotted as grand median and analyzed with one-way ANOVA. \*\**P* < 0.01, \*\*\**P* < 0.001, \*\*\*\**P* < 0.0001. Colonies were derived from HD 8, HD 16, HD 17 and HD 28. (J) Accessible reads from ATAC-seq analysis with an intact TGN7-9WGATAR motif or WGATAR motif following editing with sg1617 at +58 enhancer, following editing with sg1450 at +55 enhancer, or following editing with sg1617+sg1450 at the programmed deletion junction, +58 inversion junction and +55 inversion junction, as compared to non-targeted controls or editing with neutral locus targeting Safe sgRNA. Data are plotted as mean±SEM, n = 3 healthy donors (HD 1, HD 2 and HD 3).

We compared the editing outcomes for two additional pairs of guide RNAs targeting the +55 and +58 enhancers. We used the sg1617 adjacent sgRNA #1618, aka sg1618, targeting the +58 enhancer with sg1450 targeting the +55 enhancer, or sg1617 with the sg1450 adjacent sgRNA #1449, aka sg1449, targeting the +55 enhancer and found a similar distribution of editing outcomes as observed with the sg1617 and sg1450 pair. Gene edits persisted at high levels in engrafting cells, for sg1617+sg1449 and sg1618+sg1450 showing 86.5% and 99.5% overall editing, 35% and 31.7% programmed 3.1 kb deletions, 43.2% and 27% programmed 3.1 kb inversions, and 18% and 30.2% programmed short indels, respectively (**Figures 2B**-**2E**). We observed similar overall human chimerism in unedited and combined +58 and +55 edited recipients (mock 91.8**%**, sg1617+sg1449 90.3%, sg1618+sg1450 94.3%) (**Figure S3A**).

Multilineage repopulation was intact and *in vivo* HbF induction was robust following combined +58 and +55 *BCL11A* enhancer editing with these alternative sgRNA pairs (**Figures S3B**-**S3E**). Overall these results show that combined *BCL11A* enhancer editing with dual cleavage at the +58 and +55 core regulatory sequences with a variety of sgRNAs produces highly efficient gene editing, human chimerism and HbF induction in long-term engrafting cells.

As an independent method to determine editing outcomes, we genotyped single cell derived erythroid colonies after gene editing. We performed 5 different PCR reactions per colony to amplify each target site as well as programmed 3.1 kb deletions and inversions and performed Sanger sequencing to determine the sequence at each potential edit junction. We found that 45.6% of alleles had programmed 3.1 kb deletions, 27.2% had programmed 3.1 kb inversions, and 27.2% had indels following sg1617+sg1450 editing (**Figure 2F**). Similar results were observed for other combined editing sgRNA pairs, with programmed 3.1 kb deletions and programmed 3.1 kb inversions together constituting at least half of the repair alleles (**Figures S3F**-**S3H**). These results of individual colony analyses agreed with the ddPCR results in terms of major editing allelic outcomes.

By sequencing colonies, we were able to classify the nature of residual motif sequences at the +58 and +55 enhancers. For each colony, we classified the TGN7-9WGATAR half E-box/GATA motifs as disrupted, partially disrupted (TGN7-9 lost but WGATAR retained), or intact. We found that 58.8% of colonies had all 4 motifs disrupted and an additional 19.1% had 3 motifs disrupted following sg1617+sg1450 editing (**Figure 2G**). After sg1618+sg1450 editing, 20.3% of colonies had all 4 motifs disrupted and an additional 42.2% of colonies had 3 motifs disrupted (**Figure S3I**). Although precise inversion junctions disrupted the TGN7-9WGATAR motif, we found that a +1(A) insertion at the +58 inversion junction or a -1 deletion at the +55 inversion junction could restore TGN7-9WGATAR motifs (**Figure 2H**). Therefore a minority of the repair alleles could preserve the motif. Additionally for sg1618+sg1450 editing, some indels and inversion junctions only partially disrupt the TGN7-9WGATAR motifs and for sg1617+sg1449, some inversion junctions preserve the motifs (**Figures S3J** and **S3K**). To correlate the genotype to the HbF expression level, we used some cells from each colony for genotyping and others for hemoglobin HPLC. We found that colonies with all 4 TGN7-9WGATAR motifs disrupted showed the greatest induction of HbF (**Figure 2I**). We observed a dose-dependent relationship where colonies with fewer motifs disrupted displayed lower HbF levels. Clones with partially disrupted motifs (TGN7-9 lost but WGATAR retained) showed similarly high HbF levels as those with fully disrupted motifs, suggesting that loss of the half E-box sequence could potently impair enhancer function. Median HbF level was 66.2% for 57 clones with 0 of 4 intact TGN7-9WGATAR motifs, median HbF level was 69.2% for 38 clones with 1 or 2 WGATAR motifs intact and the other motifs fully disrupted, median HbF level was 50.8% for 24 clones with 1 of 4 intact TGN7- 9WGATAR motifs, and median HbF level was 28.9% for 27 clones with 2 or 3 of 4 intact TGN7- 9WGATAR motifs, as compared to median HbF level of 6.4% for 17 unedited clones (**Figure 2I**).

As an independent approach, we analyzed the sequences found in accessible chromatin by ATAC-seq following gene editing to evaluate the presence of core motifs. Editing the +58 enhancer with sg1617 reduced the number of reads aligning to this region, with residual reads almost all representing alleles with the TGN7-9WGATAR motif intact. Editing the +55 enhancer with sg1450 reduced reads aligning to this region, although most of the residual reads showed the motif was disrupted, suggesting other TF binding events within this enhancer could preserve chromatin accessibility in the absence of the core half E-box/GATA binding site. Combined editing of the +58 and +55 enhancers with sg1617+sg1450 reduced reads aligning to both the +58 and +55 enhancers, with nearly all residual aligning reads showing TGN7-9WGATAR motifs intact, suggesting chromatin accessibility at these enhancers was abolished in the absence of both core motifs (**Figure 2J**). Together these results suggest that disruption of core half E-box/GATA motifs at the +58 and +55 enhancers accounts for the heightened HbF induction potential of the combined enhancer editing approach.

### Specificity of combined enhancer editing

Given their potential clinical relevance, we next conducted extensive off-target analysis of sg1617, sg1618, sg1450, and sg1449. Previously, we had evaluated 24 candidate off-target sites for SpCas9 editing with sg1617 based on CIRCLE-seq, an empiric *in vitro* genomic DNA-based assay, and *in silico* homology-based CRISPOR analysis and found no off-target editing above a threshold of 0.1% allele frequency^28^. An independent analysis using GUIDE-seq, an empiric cell-based assay, and *in silico* homology analysis, discovered no off-targets for sg1617 editing in CD34+ HSPCs at 223 candidate sites, detected to a threshold of 0.2% allele frequency^6^.

We recently reported *in silico* nomination of homologous off-target sites for sg1617 incorporating human genetic variant data using the CRISPRme tool, which revealed an off-target site due to the minor allele of the SNP rs114518452^44^. The minor allele of this SNP, which is common in African ancestry populations, creates an NGG PAM sequence for an off-target site with 3 mismatches. We found that editing with a high fidelity SpCas9 variant (R691A, aka HiFi^45^) reduced off-target indels in heterozygous rs114518452-G/C donor CD34+ HSPCs from 4.8 ± 0.5% to 0.1 ± 0.1% and rendered pericentric inversions undetectable with 82.3 ± 1.6% on-target editing^44^. Based on this observation, we utilized 3xNLS-HiFi-SpCas9 for further studies to maximize editing specificity. Combined enhancer editing with 3xNLS-HiFi-SpCas9 and sg1617+sg1450 yielded comparable human chimerism and similar multilineage repopulation as mock edited controls, preserved high levels of edits in engrafting cells, and achieved potent *in vivo* HbF induction (**Figures S4A**-**S4F**).

For comprehensive off-target analysis of sg1617, sg1618, sg1450, and sg1449, we used three orthogonal methods for off-target site nomination: 1) *in silico* search for homologous genomic off-targets by CRISPRme; 2) *in vitro* evaluation by ONE-seq^46, 47^ oligonucleotide library based search; and 3) cell based evaluation by GUIDE-seq, conducted in CD34+ HSPCs (**Figures S5A**-**S5D**). In total, we nominated 315 candidate off-target sites for sg1617, 207 candidate off-target sites for sg1618, 213 candidate off-target sites for sg1450 and 343 candidate off-target sites for sg1449. We attempted pooled amplicon sequencing (rhAmpSeq^48^) of these 1078 sites following combined +58 and +55 editing from at least 3 individual CD34+ HSPC donors and achieved sufficient sequencing depth based on pre-specified criteria for all but 19 of the candidate off-target sites (**Figure S5E** and **S5F**). With the exception of minor allele rs114518452-C specific indels at the variant-associated off-target site previously described^44^, we found no significant differences in off-target indels between unedited and edited cells at an allele frequency of 0.1% for any of the 1059 tested candidate off-target sites (**Figure 3**).

**Figure 3.**
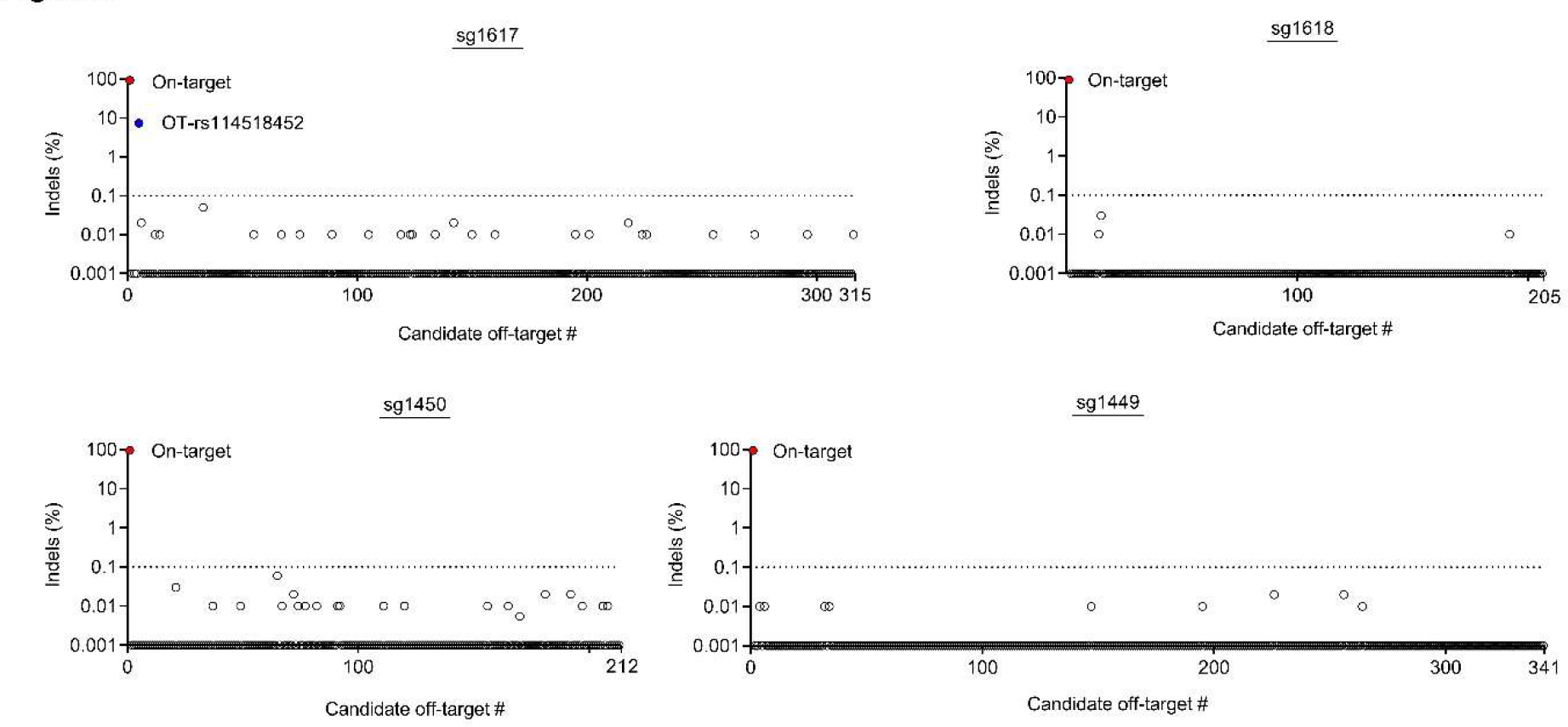
Off-target analysis of combined *BCL11A* +58 and +55 enhancer editing Candidate off-target sites nominated by CRISPRme, GUIDE-seq, and ONE-seq were subject to pooled amplicon sequencing in human CD34+ HSPCs following RNP editing targeting the *BCL11A* +58 and +55 enhancers. The mean difference between indels in edited and control samples is shown for 315 candidate off-target sites for sg1617 in HPSCs from 8 healthy donors (HD 10, HD 11, HD 15, HD 22-24, HD 29 and HD 30) edited with sg1617+sg1450 (including 4 variant-associated off-target sites with a single informative donor); 205 candidate off-target sites for sg1618 in HSPCs from 3 healthy donors (HD 7, HD 13 and HD 14) edited with sg1618+sg1450 (including 2 variant-associated off-target sites with a single informative donor); 212 candidate off-target sites for sg1450 in HSPCs from 8 healthy donors (HD 10, HD 11, HD 15, HD 22-24, HD 29 and HD 30) edited with sg1617+sg1450 (including 3 variant-associated off-target sites from a single informative donor); and 341 candidate off-target sites for sg1449 in HSPCs from 2 healthy donors and 1 sickle cell patient (HD 28, HD 31 and SCD 3) edited with sg1618+sg1449 (including 5 variant-associated off-target sites with a single informative donor). On-target editing measured by drop-off ddPCR assay. Off-target editing is evaluated by amplicon sequencing analysis. The threshold of indel frequency for true-positive significant off-target editing is set as 0.1% increase in the edited sample as compared to in the paired control sample in the mean of informative donors (dotted line).

### *Ex vivo* editing conditions determine genotoxicity

Recent reports have indicated several potentially unwanted on-target allelic outcomes after genome editing by endonucleases like Cas9. DSB repair can produce kilobase scale deletions based on extended unidirectional resection^21^. If a DSB is not repaired prior to mitosis, this can result in missegregation of the acentric chromosome fragment leading to micronuclei formation, which in turn could lead to chromothripsis^22^, and reciprocal unbalanced retention of the centric chromosome fragment^49^. Finally, recombination of edited homologous chromosomes after DNA replication could lead to uniparental disomy and copy-neutral loss-of-heterozygosity^50, 51^. We noted that each of these editing outcomes is predicted to be specific to cycling cells, which have active end-resection machineries and are susceptible to chromosome missegregation or rearrangement before mitosis is completed. We measured the cell cycle status of G-CSF mobilized PB CD34+ HSPCs from three healthy donors and plerixafor-mobilized PB CD34+ HSPCs from a SCD patient immediately after collection, with 24 hours or 48 hours of culture with cytokines (SCF, TPO, and FLT3-L each at 100 ng/ml). Flow cytometry showed that during the first 24 hours of culture the cells mainly gained biomass as indicated by increased median forward scatter and a shift from G0 towards G1 without entering S-phase (**Figures 4A** and **S6A**- **S6B**). We found 99.9±0.2% of cells to be in G0 or G1 at 0 hours, 95.1**%**±1.7% in G0 or G1 at 24 hours, and 76.1**%**±2.4% in G0 or G1 at 48 hours (**Figure 4B**).

**Figure 4.**
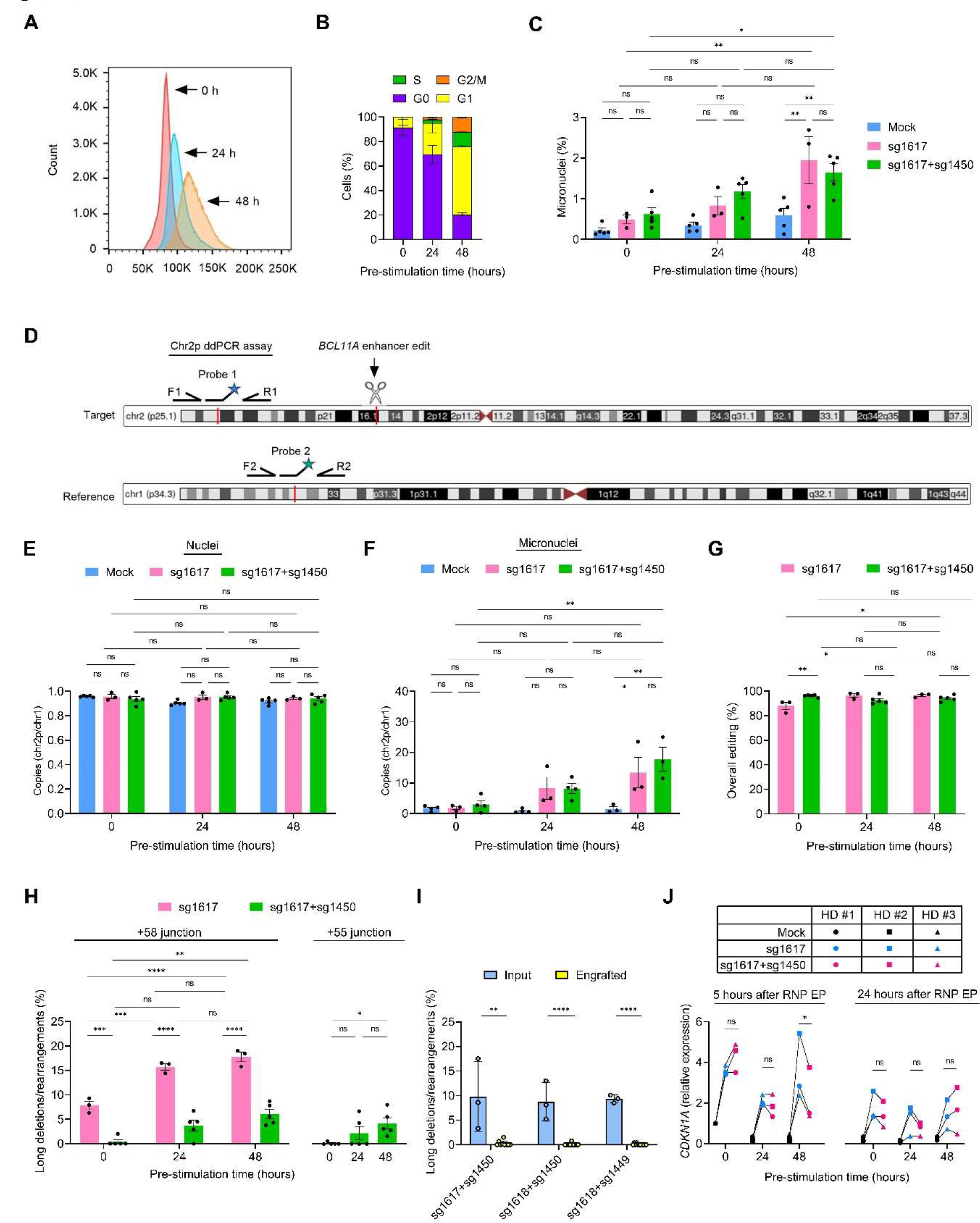
Gene editing HSPCs without *ex vivo* culture evades genotoxicity. (A) Cell size by forward scatter without (0 h) and with 24 h and 48 h of cytokine culture. (B) Fractions of cells in G0, G1, S and G2/M immediately after thawing or selection, 24 h and 48 h of pre-stimulation culture from three cryopreserved healthy donors mobilized with G-CSF and one fresh SCD patient donor mobilized with plerixafor (HD 2, HD 3, HD 31 and SCD 3) . Data are plotted as mean ± SEM. (C) Micronucleus analysis by In Vitro Micro Kit 48 h after RNP electroporation in CD34+ HSPCs without and with 24 h and 48 h of cytokines culture. Data are plotted as mean ± SEM and analyzed with one-way ANOVA. ns: nonsignificant, \**P* < 0.5, \*\**P* < 0.01. n = 3-5 healthy donors (HD 2, HD3 and HD 32-34). (D) Design of ddPCR assays recognizing chr2p (telomeric to the *BCL11A* cleavage site) and chr1 as a reference autosome. (E) Enrichment of chr2p in sorted nuclei by ddPCR assay. Data are plotted as mean ± SEM and analyzed with one-way ANOVA. ns: nonsignificant. n = 3-5 healthy donors (HD 2, HD3 and HD 32-34). (F) Enrichment of chr2p in sorted micronuclei by ddPCR assay. Data are plotted as mean ± SEM and analyzed with one-way ANOVA. ns: nonsignificant, \**P* < 0.5, \*\**P* < 0.01. n = 3-4 healthy donors (HD 2, HD3 and HD 32-34). (G) Overall editing by +58 drop-off ddPCR assay. Data are plotted as mean ± SEM and analyzed with one-way ANOVA. ns: nonsignificant, \**P* < 0.5, \*\**P* < 0.01. n = 3-5 healthy donors (HD 2, HD3 and HD 32-34). (H) Frequency of long deletions/rearrangements (not including the templated 3 kb deletion/inversion expected for double editing) was calculated as overall edit - deletion - inversion - small indels from four ddPCR assays. *Left* was measured by +58 drop-off and +58 offset assays. *Right* was measured by +55 drop-off and +55 offset assays. Data are plotted as mean ± SEM and analyzed with one-way ANOVA. ns: nonsignificant, \**P* < 0.05, \*\**P* < 0.01, \*\*\**P* < 0.001, \*\*\*\**P* < 0.0001; n = 3-5 healthy donors (HD 2, HD3 and HD 32-34). (I) Frequency of long deletions/rearrangements (not including the templated 3 kb deletion/inversion expected for double editing) in engrafting cells as compared to input cells. Data are plotted as median with range. n = 3 donors for input (SCD 3, HD 29 and HD 35 for sg1617+sg1450 and sg1618+sg1449. HD 7, HD 13 and HD 14 for sg1618+sg1450), n = 9-12 primary recipients for engrafted. (J) *CDKN1A* expression, by RT-qPCR, in CD34+ HSPCs 5 (left) and 24 (right) hours after RNP electroporation from three healthy donors. Relative expression normalized to control sample 5 hours of pre-stimulation. Data are analyzed with the unpaired two-tailed Student’s *t*-test. ns: nonsignificant. \**P* < 0.05. n = 3 healthy donors (HD 1, HD 2 and HD 3).

Most clinical protocols for autologous HSC gene therapy rely on pre-stimulation culture before gene modification as well as post-modification culture, such that the total duration in cytokine culture may be 48-72 hours or longer. We hypothesized that *ex vivo* cytokine stimulation might be dispensable for therapeutic gene editing approaches that are active in non-dividing cells, such as for NHEJ-based *BCL11A* enhancer disruption. In addition, editing HSPCs without *ex vivo* cytokine culture might protect cells from unwanted on-target genome editing outcomes. To test this, we treated CD34+ HSPCs with 48 hours of cytokine culture prior to 3xNLS-HiFi- SpCas9:sgRNA electroporation. In addition to sg1617, we used a non-targeting (NT) sgRNA as a negative control and a low-specificity (LS) sgRNA as a positive control, previously observed to have 2 off-target sites with similar edit frequencies as the target site^52^. After 48 additional hours in culture following electroporation, we evaluated for micronuclei using a flow cytometry assay that relies on sequential labeling to quantify micronuclei as membrane-bounded DNA structures from living cells with subnuclear content relative to nuclei. We observed 0.3%±0.12% micronuclei for sgNT, 1.3%±0.04% micronuclei for sg1617 and 6.7% ±2.5% micronuclei for sgLS edited cells (**Figures S6C** and **S6D**). p21 (*CDKN1A*) expression after sgLS editing was significantly higher than editing with sg1617 and sgNT (**Figure S6E**). We then tested CD34+ HSPCs with 0, 24, or 48 hours of cytokine culture before performing 3xNLS-HiFi- SpCas9:sgRNA electroporation. Following sg1617 alone or sg1617+sg1450 editing, we found that the micronucleus frequency was increased in HSPCs electroporated with Cas9 RNP following 48 hours of cytokine stimulation relative to both unedited HSPCs and HSPCs edited without preceding cytokine stimulation (**Figures 4C** and **S6F**). To evaluate if the micronuclei were indeed acentric fragments of chr2 due to *BCL11A* editing and chromosome missegregation, we performed ddPCR on sorted nuclei and micronuclei using assays specific to distal chromosome 2p and to chr1 as a reference autosome (**Figure 4D**). While we observed no enrichment of chr2p relative to the reference autosome in nuclei, we found 4.4-fold and 7.2-fold enrichment of chr2p in micronuclei following sg1617 editing and 2.7-fold and 6.0-fold enrichment of chr2p in micronuclei following sg1617+sg1450 editing after 24 and 48 hours of pre-stimulation cytokine culture. In contrast, we did not observe any enrichment of chr2p in sorted micronuclei in cells edited with either sg1617 alone or sg1617+sg1450 without pre-stimulation culture, suggesting that chromosome fragment missegregation and unwanted downstream consequences could be mitigated by editing quiescent HSCs (**Figures 4E** and **4F**).

Another unintended DSB repair outcome of gene editing can be long deletions or rearrangements. Based on the design of our ddPCR assays, long deletions or rearrangements could be inferred by the difference between overall edits and the sum of programmed deletions, programmed inversions, and short indels (**Figure S6G**). We observed that the frequency of long deletions or rearrangements was reduced in cells edited by sg1617 without cytokine culture as compared to those with 24 or 48 hours of pre-stimulation cytokine culture (**Figures 4G-4H and S6H**). The frequency of long deletions or rearrangements (excluding the programmed 3.1 kb deletion or inversion) was similarly reduced in cells edited with sg1617+sg1450 without as compared to with pre-stimulation cytokine culture, with an even more profound reduction of long deletions or rearrangements, typically to frequencies below the limit of detection (**Figures 4G- 4H** and **S6I-S6J**). Furthermore, we observed a reduced frequency of long deletions or rearrangements in engrafting cells in xenograft recipient bone marrow as compared to input cells following editing of sg1617+sg1450, sg1618+sg1450 and sg1618+sg1449 (**Figure 4I**), consistent with prior observations that gene edit distribution in engrafting HSCs may differ from that in CD34+ HSPC input cell products^28, 53^. Finally we evaluated the impact of combined enhancer editing and time of pre-stimulation culture on the p53-dependent DNA damage response by measuring p21 mRNA level. Combined +58 and +55 *BCL11A* enhancer editing with sg1617+sg1450 did not increase p21 expression in HSPCs as compared to sg1617 editing alone, independent of time in pre-stimulation culture, either at 5 or 24 hours post-electroporation (**Figure 4J**). We did observe the basal level of p21 in the unedited cells was higher without as compared to with 24-48 hours of prestimulation culture (**Figure 4J**).

We next asked if pre-stimulation culture was necessary to promote hematopoietic engraftment of gene edited cells. We compared our usual protocol of 24 hours of pre-stimulation cytokine culture, RNP electroporation, and 24 hours of post-electroporation cytokine culture, followed by cryopreservation, to a protocol without any cytokine culture. We used fresh plerixafor-mobilized PB CD34+ HSPCs from a SCD patient and fresh G-CSF mobilized PB CD34+ HSPCs from a healthy donor and electroporated immediately after selection, and then cryopreserved 15 minutes after electroporation, without any exposure to exogenous cytokines. Additionally we tested cryopreserved G-CSF mobilized PB CD34+ HSPCs from two healthy donors, electroporated immediately after thawing, and then cryopreserved 15 minutes later without cytokine culture. We compared non-electroporated control cells and cells edited at the *BCL11A* enhancer with sg1617+sg1450 or sg1618+sg1449 along with 3xNLS-HiFi-SpCas9. After 3 weeks, cells were thawed and immediately infused to NBSGW mice. A small aliquot of cells was cultured for 5 days to estimate gene edits in the input cell product. Overall gene edit frequencies were similar when comparing editing *ex vivo* without and with pre-stimulation culture, with a similar distribution of programmed deletions, programmed inversions, and small indels across conditions (**Figures 5A** and **5B**). Editing with sg1617+sg1450 or sg1618+sg1449 resulted in robust HbF induction with and without pre-stimulation culture (**Figure 5C**). After 16 weeks, BM was collected from the mice. Overall human engraftment was similar for all groups, ranging from 91.9% to 95.6% (**Figure 5D**). Gene edit frequency was preserved for recipients of HSPCs without or with pre-stimulation culture (73.7% for unstimulated for sg1618+sg1449, 79% for 24 hour pre-stimulated for sg1618+sg1449, 65.7% for unstimulated for sg1617+sg1450, 86.8% for 24 hour pre-stimulated for sg1617+sg1450) (**Figure 5E**). We observed similar multilineage repopulation of BM B-lymphoid cells, monocytes, granulocytes, and HSPCs, with a trend towards modest reduction of erythroid contribution, yet no difference in enucleation frequency after gene editing (**Figures S7A**-**S7F**). HbF levels of BM erythroid cells were potently induced, from 1.0% and 0.9% in unedited to 33.9% and 30.3% in sg1618+sg1449 edited and 29.2% and 32.9% in sg1617+sg1450 edited recipients without and with pre-stimulation culture (**Figure 5F**). Similar increases were seen for F-cells in the BM of recipients of cell products edited without or with cytokine culture (**Figure S7G**). Overall these results suggest that gene editing of non-dividing HSPCs without cytokine culture is compatible with potent gene modification and therapeutic potential and may bypass risk of genotoxicity.

**Figure 5.**
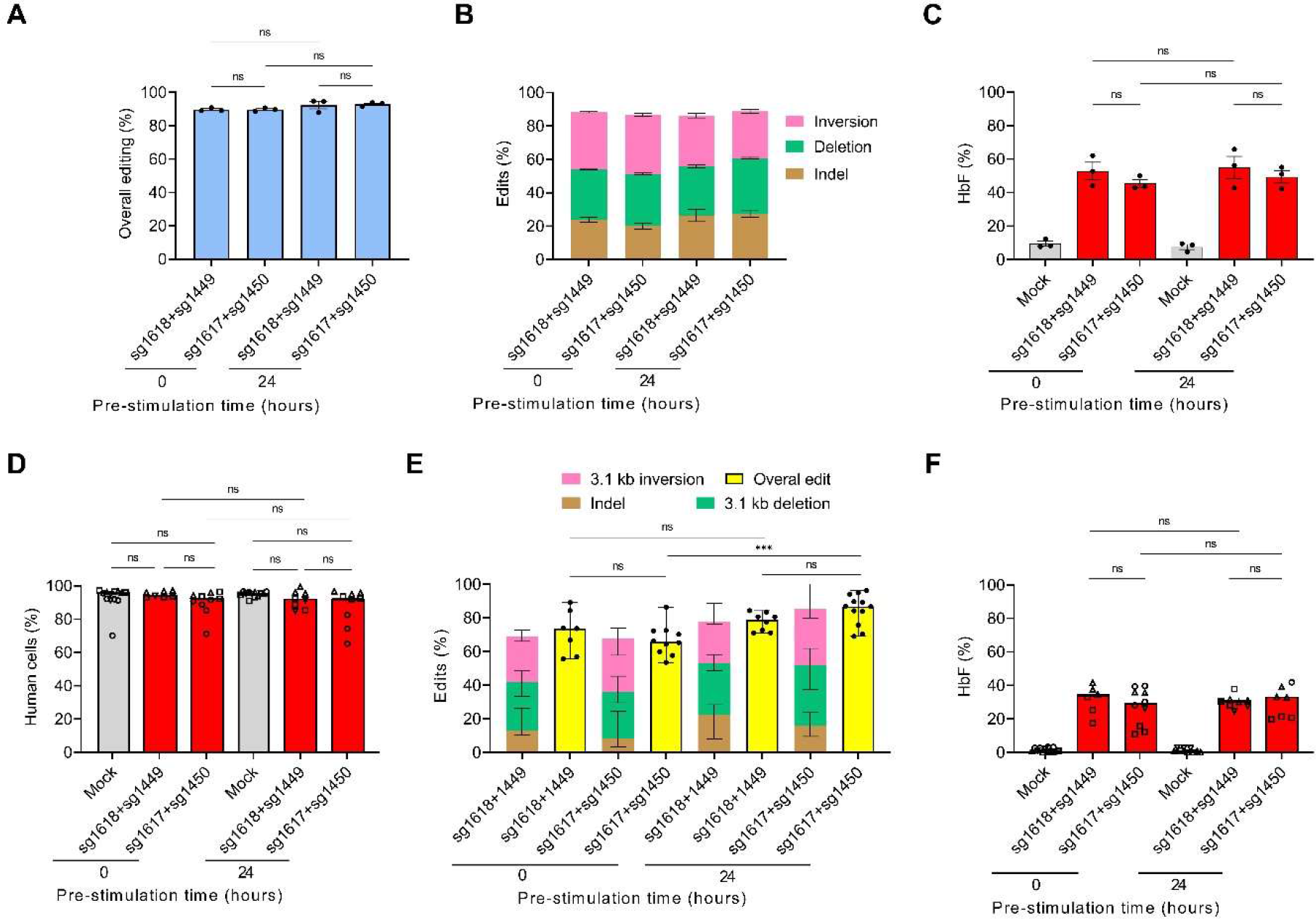
*Ex vivo* editing without cytokine culture produces potent engraftment, preserves gene edits, and reactivates HbF (A) Overall editing in *ex vivo* edited HSPCs by +58 drop-off ddPCR assay. Data are plotted as mean ± SEM and analyzed with one-way ANOVA. ns: nonsignificant. n = 2 healthy donors and 1 SCD donor (HD 28, HD 29 and SCD 3). (B) Frequencies of deletion, inversion and indels measured by ddPCR assays after *ex vivo* editing without and with 24 h of pre-stimulation culture for sg1618+sg1449 and sg1617+sg1450 editing. Data are plotted as mean ± SEM, n = 2 healthy donors and 1 SCD donor (HD 28, HD 29 and SCD 3). (C) HbF levels by HPLC in *in vitro* erythroid differentiated progeny. Data are plotted as mean ± SEM, n = 2 healthy donors and 1 SCD donor (HD 28, HD 29 and SCD 3). (D) Human chimerism 16 weeks after infusion of human CD34+ HSPCs without and with 24 hours of cytokine culture. Data are plotted as grand median and analyzed with one-way ANOVA. ns: nonsignificant. n = 7-13 primary recipients infused with edited HSPCs with or without cytokine culture from 3 healthy donors and one SCD donor (HD 28, HD 29, HD 35 and SCD3). (E) Frequencies of indel, 3.1 kb deletion, 3.1 kb inversion and overall edit by ddPCR assays in engrafted BM cells as compared to input cells. Data are plotted as median with range and analyzed with the one-way ANOVA. ns: nonsignificant, \*\*\**P* < 0.001. n = 7-12 primary recipients infused with edited HSPCs with or without cytokine culture from 3 healthy donors and 1 SCD donor (HD 28, HD 29, HD 35 and SCD3). (F) HbF levels by HPLC in engrafted erythroid cells. Data are plotted as grand median and analyzed with one-way ANOVA. ns: nonsignificant. n = 6-13 primary recipients infused with HSPCs from HD 29 and SCD 3.

## DISCUSSION

Somatic genome editing of tissue stem cells by programmable nucleases promises to correct mutations and confer beneficial properties to treat a wide range of diseases. Like most adult tissue stem cells, HSCs are largely quiescent^54, 55^. Therefore, gene repair pathways active in nondividing cells, such as NHEJ, could be harnessed to achieve therapeutic edits^56, 57^.

In this work, we demonstrate that NHEJ-based combined disruption of the two major *BCL11A* erythroid enhancers +58 and +55 produces superior HbF induction as compared to competing approaches in clinical development such as disruption of the +58 enhancer alone^6, 30^ (NCT03745287, NCT03655678, NCT04211480), disruption of *HBG1/2* promoter repressive sequences such as the -115 BCL11A binding site (NCT04853576, NCT05456880), and erythroid-restricted knockdown of *BCL11A*^7^ (NCT05353647). Greater HbF induction could be clinically meaningful to reduce residual deoxyhemoglobin polymerization and hemolysis in SCD and to limit residual ineffective erythropoiesis in β-thalassemia.

The basis for the enhanced potency of combined disruption of the +58 and +55 enhancers is loss of two core half E-box/GATA motifs, one each at the +58 and +55 enhancer. In the absence of these motifs, chromatin accessibility is almost completely abrogated at these enhancers as well as the neighboring +62 enhancer. These results support, at least in the case of the clustered erythroid enhancers at *BCL11A* intron 2, a hierarchical organization of motifs within regulatory elements, with individual core TF binding sites being required for enhancer function.

We report extensive off-target analysis for 4 SpCas9 NGG PAM restricted sgRNAs targeting these core enhancer TF binding sites. Aside from an alternative allele-specific off-target site that could be mitigated by high-fidelity SpCas9 or avoided by gRNA or donor selection^44^, we did not detect off-target editing in edited HSPCs at a threshold of 0.1% allele frequency at over 1000 candidate sites, consistent with the excellent specificity associated with transient delivery of high-fidelity Cas9 RNP^45, 58^.

By design, SpCas9-based gene editing introduces DSBs to target sequences. DSBs trigger DNA damage responses and can lead to heterogeneous outcomes, including long deletions and rearrangements^21^, copy neutral loss-of-heterozygosity^50, 51, 59^, and micronuclei, copy number loss, and chromothripsis^22^ which may contribute to unintended outcomes in therapeutic gene editing. The frequency and impact of these allelic events in HSCs after gene editing remains incompletely described. Here we show that paired DSBs targeting the *BCL11A* erythroid enhancers lead to similar p21 induction as single DSBs targeting a single enhancer. We hypothesize that the increased burden of multiple DSBs is offset by the generation of frequent programmed 3.1 kb deletions and inversions that prevent cycles of perfect repair and re-cleavage. Furthermore we find that unintended outcomes like micronuclei are no more frequent and long deletions even less frequent following paired as compared to single programmed DSBs at the *BCL11A* enhancers. Both micronuclei and long deletions are expected to occur in dividing cells (in M phase for micronuclei^60, 61^ and S/G2 for extensive end resection^56^). Induction of cell cycle through *ex vivo* cytokine stimulation and extended *in vitro* culture is also negatively correlated with HSC engraftment potential^54, 62, 63^.

Therefore, we reasoned that editing nondividing HSPCs without *ex vivo* cytokine culture could achieve frequent NHEJ-mediated intended allelic outcomes, preserve HSC engraftment function, and bypass unintended genotoxicities associated with DSBs in dividing cells. Indeed gene edits like MMEJ and HDR that are depleted from immunophenotypic HSCs and G0 cells within CD34+ HSPCs are selectively reduced in engrafting cells, suggesting that nondividing HSCs possess unique DNA damage repair preferences as compared to rapidly dividing progenitors^20, 28, 30, 53, 64–66^. We observed that mPB HSPCs are quiescent after CD34 selection and only enter G1 and then S phase after 24-48 hours of *ex vivo* cytokine culture. Merely delivering SpCas9 RNPs for combined *BCL11A* enhancer editing to HSPCs without cytokine culture was sufficient to prevent generation of micronuclei and long deletions. Importantly, edited CD34+ HSPCs without cytokine culture were competent for long-term multi-lineage repopulation, efficient gene editing, and potent HbF induction, indicating that *ex vivo* culture may be dispensable for therapeutic gene editing.

In contrast to intended NHEJ therapeutic edits, the need for protocols to promote S/G2 phase for gene repair via HDR^67^ and co-delivery of high concentration DNA template may diminish the repopulation potential of HSCs, which may be related to challenges in achieving hematopoietic recovery following *ex vivo* HDR editing protocols^68^. Given the heterogeneity of gene repair outcomes, which may be modified by cell intrinsic and extrinsic factors, we expect that input cell product characterization may incompletely reflect edit distribution in HSCs. Therefore we would encourage all clinical trials of therapeutic HSC editing to correlate editing procedures with allelic outcomes in engrafting cells.

Current gene therapy approaches for blood disorders rely on *ex vivo* cell manipulation prior to myeloablative conditioning, an approach that is complex, expensive and toxic, and cannot easily scale to the global problem of congenital blood disorders let alone diseases like HIV, autoimmunity, and cancer that could benefit from such therapies. *In vivo* delivery of gene editing reagents to HSCs would appear to be a promising avenue for simplifying and improving access to therapeutic gene editing, yet the feasibility and safety of targeting HSC *in situ* is unknown.

These studies offer encouragement that editing quiescent HSCs, as would be needed for *in vivo* delivery, may be feasible and could in fact protect against genotoxicity.

## METHODS

### Cell culture

Cryopreserved human CD34^+^ HSPCs from mobilized peripheral blood of deidentified healthy donors were obtained from Fred Hutchinson Cancer Research Center, Seattle, Washington. Fresh mobilized peripheral blood from healthy donors (G-CSF and G-CSF+plerixafor) (see donor information in **Table S1**) were obtained from either Hemacare Corp (Northbridge, California) or Miltenyi Biotec (Auburn, California). Plerixafor mobilized peripheral blood from sickle cell disease patients were procured under protocols approved by the IRB of Boston Children’s Hospital. CD34^+^ HSPCs were isolated using the CliniMACS® CD34 reagent (Miltenyi, cat# 130-017-501) and CliniMACS® LS tubing set (Miltenyi, cat# 170-076-651). CD34^+^ HSPCs were thawed and cultured into Stem Cell Growth Medium (SCGM), GMP grade (CellGenix, cat#, 20806-0500) supplemented with 100 ng ml^-1^ Preclinical Thrombopoietin (TPO) (CellGenix, cat# 1417-050), 100 ng ml^-1^ Preclinical Stem Cell Growth Factor (SCF) (CellGenix, cat# 1418- 050) and 100 ng ml^-1^ Preclinical FMS-like Tyrosine Kinase 3 Ligand (FLT3L) (CellGenix, cat# 1415-050). HSPCs were electroporated with 3xNLS-SpCas9:sgRNA or 3xNLS-HiFi- SpCas9:sgRNA RNP immediately, 24 hours or 48 hours after thawing or selection. HSPCs were cultured in SCGM medium with cytokines for another 24 hours after electroporation unless otherwise stated. For *in vitro* erythroid differentiation experiments, 24 hours after electroporation, HSPCs were transferred into erythroid differentiation medium (EDM) consisting of IMDM (GibcoTM, 12440061) supplemented with 330 μg/ml holo-human transferrin (Sigma, T0665-1G), 10 μg/ml recombinant human insulin (Sigma, 19278-5ML), 2 IU/ml heparin (Sigma, 19278-5ML), 5% human solvent detergent pooled plasma AB (Rhode Island Blood Center), 3 IU/ml erythropoietin (AMGEN, 55513-144-10). During days 0-7 of culture in EDM-1, EDM was further supplemented with 10^-6^ M hydrocortisone (Sigma-Aldrich, H0135), 100 ng/ml human SCF (CellGenix, 1418-050) and 5 ng/ml of recombinant human IL-3 (PEPROTECH, 200-03). During days 7-11 of culture in EDM-2, EDM was supplemented with 100 ng/ml human SCF (CellGenix, 1418-050). During days 11-18 of culture in EDM-3, EDM had no additional supplements.

### RNP electroporation

HSPCs were electroporated using Lonza 4D nucleofector or MaxCyte GT device. For Lonza 4D small scale electroporation, 300 pmol (15 μM) of sg1617 or sg1618 (see sequences of sgRNAs in **Table S2**) was mixed with 100 pmol (5 μM) of 3xNLS-SpCas9 protein and added electroporation buffer (Lonza 4D, cat# V4XP-3032) up to 5 μl in one tube, 300 pmol (15 μM) of sg1450 or sg1449 was mixed with 100 pmol (5 μM) of 3xNLS-SpCas9 protein and added electroporation buffer up to 5 μl in another tube. RNP from two tubes was mixed after 15 min of incubation at room temperature to allow initial separate RNP complexing of each sgRNA with Cas9. 50,000-100,000 HSPCs were prepared and resuspended in 10 μl of electroporation buffer, RNP were mixed with cells and transferred to cuvette (Lonza 4D, cat# V4XP-3032).

Electroporation was performed using the program EO-100 with Lonza 4D nucleofector. For Lonza 4D large scale electroporation, 1500 pmol (15 μM) of sg1617 or sg1618 was mixed with 500 pmol (5 μM) of 3xNLS-SpCas9-SpCas9 protein and added electroporation buffer (Lonza 4D, cat# V4XP-3024) up to 25 μl in one tube, 1500 pmol (15 μM) of sg1450 or sg1449 was mixed with 500 pmol (5 μM) of 3xNLS-SpCas9 protein and added electroporation buffer up to 25 μl in another tube. RNP from two tubes was mixed 15 min of incubation at room temperature.

5,000,000 HSPCs were prepared and resuspended in 50 μl of electroporation buffer, RNP were mixed with cells and transferred to cuvette (Lonza 4D, cat# V4XP-3024). Electroporation was performed using the program EO-100 with Lonza 4D nucleofector. For Maxcyte small scale electroporation, 750 pmol (15 μM) of sg1617 or sg1618 was mixed with 250 pmol (5 μM) of 3xNLS-HiFi-SpCas9 protein and added electroporation buffer (MaxCyte® cat# EPB1) up to 12.5 μl in one tube, 750 pmol (15 μM) of sg1450 or sg1449 was mixed with 5 μM of 3xNLS-HiFi- SpCas9 protein and added electroporation buffer up to 12.5 μl in another tube. RNP from two tubes was mixed after 15 min of incubation at room temperature. 500,000-5,000,000 HSPCs were prepared and resuspended in 25 μl of electroporation buffer, RNPs were mixed with cells and transferred to OC-100 (MaxCyte® cat# GOC1). Electroporation was performed using energy HSC-3 for HSPCs after 24 h of pre-stimulation, HSC-5 for HSPCs without *ex vivo* cytokine culture.

### Lentiviral shmiR procedures

The lentiviral vector, production of virus preparations and transduction of CD34+ cells were performed as previously described^69^. Vector copy numbers in bulk cultured cells were determined on a split of cells cultured for 7 days after transduction or from freshly isolated bone marrow from humanized mice. Genomic DNA was isolated using the Qiagen DNeasy protocol (Qiagen, cat# 69506). qPCR was performed on an Applied Biosystems 7500 Real-Time PCR system in a duplex reaction using the vector specific Taqman probe FAM- AGGCTAGAAGGAGAGAGTAGGGTGC-TAMRA and primers: Forward GACTGGTGAGTACGCCAAA and Reverse: CGCTTAATACCGACGCTCTC. GTDC1 served as reference using: Probe VIC-ACGAACTTCTTGGAGTTGTTTGCT-TAMRA and primers: Forward GAAGTTCAGGTTAATTAGCTGCTG and Reverse GGCACCTTAACATTTGGTTCTG. The VCN was calculated relative to a known reference calibrator sample containing one copy of viral DNA per diploid genome.

### Transplantation

All animal experiments were approved by Boston Children’s Hospital Institutional Animal Care and Use Committee. NOD.Cg-KitW-41J Tyr + Prkdcscid Il2rgtm1Wjl (NBSGW) female mice (4-6 weeks of age) were obtained from Jackson Laboratory (stock 026622). For HPSCs without pre-stimulation, cells were cryopreserved in CryoStor™ 5 cryomedia (Biolife Solutions cat# 205102) 15 min after electroporation and after removing the electroporation buffer. For HSPCs with 24 hours of pre-stimulation, cells were cryopreserved in CryoStor™ 5 cryomedia 24 hours after electroporation. HSPCs were thawed 3 weeks after cryopreservation and infused into NBSGW mice immediately after thaw. 1,000,000 HSPCs were resuspended in 100 μl of SCGM medium without cytokines and injected into mice via retro-orbital method^70^. Bone marrow was isolated 16 weeks after transplantation for final analysis.

### Flow cytometry

Bone marrow cells were incubated with Human TruStain FcX (BioLegend, cat# 422302, RRID: AB_2818986) and TruStain FcX (anti-mouse CD16/32) (BioLegend, cat#101320, RRID: AB_1574975) for 15 min at room temperature. Antibodies including viability dye eFluor 780 (Thermo fisher, cat# 65-0865-18), mCD45_PE_eFluor 610 (Thermo Fisher Scientific, cat# 61- 0451-82, RRID: AB_2574555), hCD45_V450 (BD Biosciences cat# 560367, RRID: AB_1645573), hCD235a_FITC (BioLegend, cat# 349104, RRID: AB_10613463), hCD19_APC (BioLegend, cat# 302212, RRID:AB_314242), hCD33_PE (BioLegend, cat# 366608, RRID: AB_2566107), hCD3_PE_Cy7 (BioLegend, cat# 300420, RRID: AB_439781), hCD34_FITC (BioLegend, cat# 343504, RRID: AB_1731852), were added to the cells and incubated on ice for 30 min. Samples were acquired on a BD FACSAria II and data were analyzed with FlowJo™ software. Populations were sorted using the general lineage panel: hCD45+hCD19+ cells (human B cells), hCD45+hCD19-hCD33-hCD34+ cells (human HSPCs), mCD45-hCD45- hCD235a+ cells (human erythroid cells), and hCD45+hCD19-hCD235a-hCD34- cells (human non-B, non-erythroid, non-HSPC cells). Sorted cells were used for RNA extraction and determination of *BCL11A* expression.

### MACS human CD235a isolation

Bone marrow cells were incubated with hCD235a microbeads (Miltenyi Biotec, cat# 130-050- 501) on ice for 15 min. Cells were isolated by LS column (Miltenyi Biotec, cat# 130-042-401). hCD235a- cells were used for secondary transplantation. 20% of hCD235a+ cells were used for F cell and enucleation analysis by flow cytometry, 80% of hCD235a+ cells were used for HbF measurement by HPLC.

### F cell and enucleation analysis

Human CD235a+ cells were incubated with Hoechst 33342 (Millipore, cat# B2261) for 20 min at 37°C, fixed with 0.05% glutaraldehyde in PBS for 10 min, and permeabilized with 0.1% Triton X- 100 in PBS+BSA for 5 min. Anti-human CD235a_APC (BD Bioscience, cat# 551336, RRID: AB_398499) and anti-human HbF-FITC (Thermo Fisher Scientific, cat# MHFH01-4, RRID: AB_2539768) were added and incubated on ice for 30 min. Samples were acquired on a BD FACSAria II and data were analyzed with FlowJo™ software.

### HPLC

Hemolysates were prepared from erythroid cells after 18 days of erythroid differentiation or engrafting erythroid cells using Hemolysate reagent (Helena Laboratories cat# 5125) and analyzed with D-10 Hemoglobin Analyzer (Bio-Rad).

### Genotyping of single cell derived erythroid colonies

Single cell derived from human CD34+ HSPCs was sorted 24 hours after RNP electroporation or shmiR transduction. The cells were sorted by BCL2-ARIA II SPEC (BD Biosciences) into 150 μL of EDM-1 in 96-well round bottom plates (Thermo Fisher Scientific, cat #3799) at one cell per well. The cells were changed into 250-500 μL of EDM-2 if colonies were visible at the round bottom of plates after 7 days of EDM-1 culture. After an additional 4 days of EDM-2 culture, the cells were changed into EDM-3 at 1,000,000 cells/ml. After an additional 7 days of EDM-3 culture, half of the cells were used for hemoglobin HPLC measurement and half of the cells were used for genotyping analysis.

### NGS library preparation and analysis of edited clones

Targeted amplicons were generated using gene specific primers with partial Illumina adapter overhangs and sequenced as previously described^71^. Briefly, cell pellets were lysed and used to generate gene specific amplicons with partial Illumina adapters in PCR 1. Amplicons were indexed in PCR 2 and pooled with other amplicons to create sequence diversity. Additionally, 10% PhiX Sequencing Control V3 (Illumina) was added to the pooled amplicon library prior to running the sample on an Miseq Sequencer System (Illumina) to generate paired 2 X 250bp reads. Samples were demultiplexed using the index sequences, fastq files were generated, and NGS analysis was performed using CRIS.py^72^. Relevant sequencing primers are shown in **Table S3**. Relevant CRIS.py analysis parameters are shown in **Table S4**.

### ATAC-seq

ATAC-seq was performed as previously described^73^ from CD34+ HSPC derived erythroid precursors after 11 days of erythroid differentiation. 50,000 cells were washed with 50 μL of cold 1x DPBS buffer (Gibco, cat #14190-250) and centrifuged at 500 ×g for 5 min at 4°C. The cell pellet was resuspended in 50 μL of cold lysis buffer (10 mM Tris-HCl, pH 7.4, 10 mM NaCl (Thermo Fisher Scientific, cat# AM9760G), 3 mM MgCl2 (MilliporeSigma, cat #5985-100ML), 0.1% IGEPAL CA-630 and centrifuged at 500 ×g for 10 min at 4°C. The cell pellet immediately proceeded to transposition reaction. The nuclei was resuspended in the transposition reaction mix containing 25 μL 2x TD Buffer (Illumina, cat #FC-121-1030), 2.5 μL Tn5 Transposase (Illumina, cat #FC-121-1030), 22.5 μL Nuclease Free Water (Thermo Fisher Scientific, cat# AM9937). The transposition reaction was incubated at 37°C for 30 min. Immediately following transposition, purification was performed using Qiagen MinElute Kit (Qiagen, cat #28004) .

Transposed DNA was eluted in 10 μL Elution Buffer (10mM Tris buffer, pH 8). Transposed DNA fragments were amplified with the reaction containing: 10 μL transposed DNA, 9.7 μL Nuclease Free Water, 2.5 μL 25 μM Customized Nextera PCR Primer 1 (**Table S3**), 2.5 μL 25 μM Customized Nextera PCR Primer 2 (**Table S3**), 0.3 μL 100x SYBR Green (Invitrogen, cat #S- 7563) and 25 μL NEBNext High-Fidelity 2x PCR Master Mix (New England Labs, cat #M0541). The cycling conditions were as follows: 72°C for 5 min, 98°C for 30 sec, 4 cycles of 98°C for 10 sec, 63°C for 30 sec and 72°C for 1 min. qPCR was performed with the reaction containing 5 μL 5 cycles PCR amplified DNA, 4.44 μL Nuclease Free Water, 0.25 μL 25 μM Customized Nextera PCR Primer 1 (**Table S3**) 0.25 μL 25 μM Customized Nextera PCR Primer 2 (**Table S3**), 0.06 μL 100x SYBR Green and 5 μL NEBNext High-Fidelity 2x PCR Master Mix. The cycling conditions were as follows: 98°C for 30 sec, 19 cycles of 98°C for 10 sec, 63°C for 30 sec and 72°C for 1 min. Run the remaining 45 μL PCR reaction to the correct cycle number based on the calculation that corresponded to ¼ of maximum fluorescent intensity after setting 5000 RF threshold from Plot linear Rn vs. Cycle. Plot linear Rn vs. Cycling conditions were as follows: 98°C for 30 sec, 3-5 cycles of 98°C for 10 sec, 63°C for 30 sec and 72°C for 1 min. The library was purified using Qiagen PCR Cleanup Kit (Qiagen, cat #28104) and eluted in 20 μL elution Buffer (10mM Tris Buffer, pH 8). Library QC was performed using gel electrophoresis, library quantitation was performed using Qubit and Tapestation and the library was sequenced by NovaSeq. The ATAC-seq full genome alignment, processing and quality control were obtained by using the ATAC-seq ENCODE pipeline^74^ using the hg38 human genome as a reference for alignment. The resulting alignment tracks were visualized in the Integrative Genomics Viewer (IGV).

For the quantification of motifs at ATAC-seq junctions, we created multiple reference sequences based on the unedited alleles at the +58 and +55 enhancers, with 20 nt flanks extending from the predicted cleavage position, as well as predicted perfect 3108 nt deletion junction and perfect +58:+55 and +55:+58 inversion junctions again with 20 nt flanks extending from the predicted cleavage position (**Table S4**). The ATAC-seq raw reads (FASTQ files) were aligned against these generated reference edit sites using Bowtie2 (v2.3.4.3) with the following parameters: “*bowtie2 -x edit_custom_ref --seed 2248 -L 5 --sam-no-qname-trunc --local*” to allow for alignments that accommodate indels. The post alignment reads were filtered with bamtools *view (*v2.5.1) to remove unaligned reads, while samtools *view* (v0.1.19) was employed to summarize aligned reads containing deletions. To avoid partial alignments, the reads not covering the start and end nucleotides from a reference region (nucleotide 1 and 40) were removed. Finally, the occurrences of the TGN7-9WGATAR motifs were computed for each edit reference for the control and edited ATAC-seq replicates. The motif matches were searched using the *Python regular expressions* limiting to three possible outcomes: Complete motif *(+) GATA1 (+) half E-box*, partial motifs *(+) GATA1 (-) half E-box* and motif absence *(-) GATA1*.

Counts for each of the ATAC-seq samples were normalized by CPM using the total reads from full genome alignment, and the motif count numbers for each replicate were summed.

### BCL11A expression levels by RT-qPCR

For *in vitro* differentiated erythroid cells, RNA isolation was performed using RNeasy Mini kit (Qiagen, cat# 74106). Reverse transcription was done using the iScript cDNA Synthesis Kit (Bio-Rad, cat# 170-8890). *BCL11A* expression was determined by ddPCR assay using ddPCR supermix (no dUTP) (BioRad, cat# 1863023) and the *BCL11A* primers: Forward: AGCTCACCAGGCACATGAAA, Reverse: CACTCGATCACTGTGCCATT and probe:

CAGGTGGGGAAGGACGTTTA. GAPDH was used as a reference gene (assay ID: Hs02758991_g1, TaqMan Gene Expression Assay, VIC (Applied Biosystems^TM^, cat #4448489). For engrafting cells, RNA isolation was performed using RNeasy Micro Kit (Qiagen, cat# 7004), and reverse transcription was done using the iScript cDNA Synthesis Kit. *BCL11A* expression was determined by SYBR Green assay using SYBR Select Master Mix (ThermoFisher, cat #4472920) and the primers for *BCL11A*: Forward: GCCCCAAACAGGAACACATA. Reverse: GGGGCATATTCTGCACTCAT. *GAPDH* was used as internal control for sorted B cells, HSPCs and non-B, non-HSPC, non-erythroid cells, Forward: ACCCAGAAGACTGTGGATGG. Reverse: TTCAGCTCAGGGATGACCTT. *Catalase* was used as internal control for sorted erythroid cells. Forward: CTTCGACCCAAGCAACATGC. Reverse: CGGTGAGTGTCAGGATAGGC.

### Droplet digital PCR

Genomic DNA was isolated using the Blood and Tissue Kit (Qiagen cat# 69506) according to the vendor’s recommendations. 50-100 ng of genomic DNA was used for each assay. ddPCR was performed using Biorad QX200 AutoDG droplet digital PCR system. ddPCR supermix (no dUTP) (BioRad, cat# 1863023) and HindIII (NEB, cat# R3104S) were added in the reaction. +58 drop-off ddPCR assay for sg1617: Forward: TCCTCTTCTACCCCAC, Reverse: GGCATCTACTCTTAGACATA, Probe: AGCCTGTGATAAAAGCAA. +58 drop-off ddPCR assay for sg1618: Forward: TCCTCTTCTACCCCAC, Reverse: GGCATCTACTCTTAGACATA, Probe: TCCTGGAGCCTGTGA. +55 drop-off ddPCR assay for sg1449 and sg1450: Forward: CTCACAGGACATGCAG, Reverse: CCCTGTGATCTTGTGG, Probe:

CCCTATCAGTGCCGAC. +58 offset ddPCR assay: Forward: TCCTCTTCTACCCCAC, Reverse: GGCATCTACTCTTAGACATA. Probe: CCCCACCCTAATCAGAG. +55 offset ddCPR assay: Forward: CTCACAGGACATGCAG, Reverse: CCCTGTGATCTTGTGG, Probe: GATGCACACCCAGGCTG. 3.1 kb deletion ddPCR assay: Forward: CCACCGATGGAGAGGTCT. Reverse: CAGCCTGGGTGTGCAT. Probe:

CCAGTCCTCTTCTACCCCACC. 3.1 kb inversion ddPCR assay: Forward: TGGCATCTACTCTTAGACATAACAC. Reverse: CAGCCTGGGTGTGCATC. Probe:

ACCAGGGTCAATACAACTTTGAAGC. Reference assay for +58 drop-off assay and +58 offset assay: *EIF2C1*, Human (Bio-Rad, 10031243, ID: dHsaCP1000002). Reference assay for 3.1 kb deletion and 3.1 kb inversion assays: *EIF2C1* refv3 (Bio-Rad, 10031279), Forward: CTAGCCATTGTGAGCTGGC, Reverse: ACCCAATACCTCATGGATGC, Probe:

CCCTGGATGTGGCCATGAGG. After droplet generation, PCR was followed the cycling conditions: 1 cycle of 95°C for 10 min, 50 cycles of 94°C for 30 sec, 56°C for 60 sec, 1 cycle of 98°C for 10 min, held at 4°C. PCR product was read by droplet reader and the results were analyzed using the QuantaSoft software. Overall editing frequency = 100 - (unedited allele frequency of edited sample by drop-off assay / unedited allele frequency of mock sample by drop-off assay) * 100. Short indel frequency = (unedited and short indel allele frequency of edited sample by offset assay / unedited and short indel allele frequency of mock sample by offset assay) * 100 - (100 - overall editing frequency by drop-off assay). Deletion frequency = (deletion allele frequency of edited sample by 3.1 kb deletion assay / (deletion allele frequency of single copy deletion clone by 3.1 kb deletion assay / 2) * 100. Inversion frequency = (inversion allele frequency of edited sample by 3.1 kb inversion assay / inversion allele frequency of single copy inversion clone by 3.1 kb inversion assay / 2) * 100. Long deletion or rearrangement frequency = Overall editing frequency by drop-off assay - small indel frequency by offset assay - deletion frequency by 3.1 kb deletion assay - inversion frequency by 3.1 kb inversion assay. Single copy deletion and inversion clones were generated by first isolating a clone with a heterozygous 11 kb deletion encompassing *BCL11A* +55 and +58 enhancers by SpCas9:sg*BCL11A*_enhancer_5’ del+sg*BCL11A*_enhancer_3’ del RNP electroporation (see sequences of sgRNAs in **Table S2**), and next by RNP electroporation with 3xNLS- SpCas9:sg1617+sg1450 and isolating subclones with single copy 3.1 kb deletion or 3.1 kb inversion respectively verified by Sanger sequencing of edit junctions.

### Sanger sequencing

*BCL11A* enhancer DHS +58 and +55 core regions were amplified with KOD Hot Start DNA Polymerase (EMD-Millipore, cat# 71086-3) and corresponding +58 Sanger primers: Forward: CACACGGCATGGCATACAAA, Reverse: CACCCTGGAAAACAGCCTGA. +55 Sanger primers: Forward: GCTGGGGTGAGTCAAAAGTC, Reverse: CATCCATCAGAGAGGCTTCC. PCR was followed the cycling conditions: 1 cycle of 95°C for 3 min, 38 cycles of 95°C for 20 sec, 60°C for 10 sec and 70°C for 10 sec, 1 cycle of 70°C for 5 min. Sanger traces were imported to TIDE CRISPR version 3.2.0 for indel measurement with 100 bp left boundary and automatically set at breaksite -10 bp as alignment window, 115-515 bp decomposition window, 40 bp indel size range and 0.001 *P*-value.

### Amplicon deep sequencing

The *BCL11A* enhancer DHS +58 and +55 target sequences were amplified with KOD Hot Start DNA Polymerase (EMD-Millipore, 71086-31) and corresponding primers: +58 Forward: AGAGAGCCTTCCGAAAGAGG, +58 Reverse: GCCAGAAAAGAGATATGGCATC; +55 Forward: ACAGTGATAACCAGCAGGGC, +55 Reverse: GATGCAATGCTTGGAGGCTG. The cycling conditions were 95◌֯ C for 3 min; 30 cycles of 95◌֯ C for 20 s, 60◌֯ C for 10 s, and 70◌֯ C for 10 s; 70◌֯ C for 5 min. Locus specific PCR product was used for indexing PCR using KOD Hot Start DNA Polymerase and TruSeq i5 and i7 indexing primers (Illumina) following the cycling conditions: 95°C for 3 min; 10 cycles of 95°C for 20 s, 60°C for 10 s, and 70°C for 10 s; 70°C for 5 min. The indexed PCR products were evaluated by Qubit dsDNA HS Assay Kit (Thermo Fisher, Q32854), TapeStation with High Sensitivity D1000 Reagents (Agilent, 5067-5585) and High Sensitivity D1000 ScreenTape (Agilent, 5067-5584) and KAPA Universal qPCR Master Mix (KAPA Biosystems, KK4824/Roche 07960140001). The products were pooled as equimolar and subjected to deep sequencing using MiniSeq (Illumina). Amplicons were sequenced using paired-end ∼150 bp reads on Illumina HiSeq, NovaSeq, and MiniSeq instruments. Reads were trimmed for adapters and quality using Trimmomatic^75^ in paired-end mode. Editing outcomes were analyzed using CRISPResso2 (v2.2.9)^76^ using the default (Cas9) parameters along with the ignore_substitutions option. CRISPRessoBatch was used for analysis.

### GUIDE-seq

GUIDE-seq_dsODN_sense: TTAATTGAGTTGTCATATGTTAATAACGGT and GUIDE-seq_dsODN_antisense: ACCGTTATTAACATATGACAACTCAATTAA were annealed following the cycling conditions: 1 cycle of 95◌֯ C for 2 min, 700 cycles of 95◌֯ C for 1 min, increment -0.1 ֯C/cycle, and held at 25◌֯ C. 5 μM of GUIDE-seq dsODN was delivered into human CD34+ HSPCs along with 30 μM sgRNA:10 μM of 3xNLS-SpCas9 RNP. The cells were harvested for genomic DNA extraction 5 days after electroporation. Library preparation utilized Tn5 transposase for adaptor incorporation as described in the GUIDE-tag protocol^77^. Tn5 enzyme (Wolfe lab, University of Massachusetts Chan Medical School) was assembled with pre-annealed oligos by incubating at room temperature for one hour. 1 µL of assembled transposome was used for tagmentation of 200 ng of genomic DNA by incubating at 55◌֯ C for 7 min. Tagmentation reaction was inactivated with 0.2% SDS (ThermoFisher, cat# 15553027).

Tagmented DNA was used for GUIDE-seq library preparation. Using 2x Platinum SuperFi PCR Master mix (ThermoFisher, cat# 12358050), dsODN_sense specific PCR product was amplified using primer: ACCGTTATTAACATATGACAACTCAATTAA. dsODN_antisense specific PCR product was amplified using primer: TTAATTGAGTTGTCATATGTTAATAACGGT. The cycling conditions were: 1 cycle of 98°C for 2 min, 15 cycles of 98°C for 10 sec, 58°C for 10 sec and 72°C for 90 sec, 1 cycle of 72°C for 5 min and held at 4°C. dsODN specific PCR products were used for indexing PCR using 2x Platinum SuperFi PCR Master mix and i5 primer: AATGATACGGCGACCACCGAGATC and i7 indexing primers (Illumina) following the cycling conditions: 1 cycle of 98°C for 2 min; 20 cycles of 98°C for 10 s, 58°C for 10 sec, and 72°C for 90 sec; 72°C for 5 min. The indexed PCR products were evaluated by Qubit dsDNA HS Assay Kit (Thermo Fisher, Q32854), TapeStation with High Sensitivity D1000 Reagents (Agilent, 5067- 5585) and High Sensitivity D1000 ScreenTape (Agilent, 5067-5584) and KAPA Universal qPCR Master Mix (KAPA Biosystems, KK4824/Roche 07960140001). The libraries were deep-sequenced as a pool using paired-end 150-bp run on an Illumina MiniSeq with the following parameters: ≥20% PhiX (or other diverse library) and 141-146|8|17|141-146 (Read1|Index1|Index2|Read2). Deep sequencing data from the GUIDE-seq experiment was demultiplexed, preprocessed, and aligned to the hg38 human reference genome with GS- preprocess^78^ followed by unique cleavage identification and off-target nomination using the Bioconductor package GUIDEseq (v1.27.8)^79^. Off-target site identification parameters were set for SpCas9 as follows: min.reads = 1, min.read.coverage = 1, max.mismatch = 10, PAM.pattern = “NNN”, includeBulge = TRUE, max.n.bulge = 2, min.umi.count = 2, upstream = 25, downstream = 25, keepPeaksInBothStrandsOnly = FALSE. To filter false positive signals due to fragile genomic sites or sequencing noise, we performed the same analysis with oligo-only samples and subtracted any signal found in the controls. This analysis provided a list of potential off-target sites ranked with regards to the number of UMIs in the peak region associated with unique coordinates (peak score), which has been correlated with the activity of SpCas9 at each sequence. As per prior standard^80^, all sites identified by GUIDE-seq in CD34+ HSPCs with at least 2 UMIs were included as candidate off-targets for amplicon sequencing to prioritize any potential targeted loci.

### ONE-seq

ONE-seq was performed as previously described^46, 47^. In brief, Cas-Designer^81^ was used to screen the human reference genome for all candidate off-target sites with up to six mismatches relative to the target DNA sequence or four mismatches and one or two DNA/RNA bulges. Potential off-target site sequences and 10 bps of flanking sequence both upstream and downstream were included in the libraries. The off-target loci were embedded in the middle of a ∼200 nt oligonucleotide, comprising constant sequence, and a unique 14 bp barcode present on both ends of each library member. Oligonucleotide libraries were synthesized by Agilent Technologies. *In vitro* Cas9 cleavage reactions were performed in 100 µL reactions with 1X Cas9 Reaction Buffer (or NEBuffer 3.1), 30 ng of amplified library, and an sgRNA:Cas9:DNA ratio of 20:10:1. 3xNLS-SpCas9 was used with sgRNA as RNP. Cleaved products were blunted at 72°C for 10 min in a 50 µL reaction containing 1X Phusion HF Buffer, 200 µM dNTPs, and 1 unit of Phusion DNA polymerase (NEB). Product purification and adapter-ligation were performed according to the manufacturer’s protocol. Gel-purified products were amplified in 50 µL reactions containing 12 µL of sample, 1X Phusion HF Buffer, 200 µM dNTPs, 0.5 µM oKP40 (see primers for ONE-seq in **Table S3**), 0.5 µM oKP101 for the protospacer side or 0.5 µM oKP154 for the PAM side, and 1 unit of Phusion DNA Polymerase (NEB). After purification with paramagnetic beads, a second PCR was performed in 50 µL reactions containing 10 µL of Protospacer or PAM side amplified selection product, 1X Phusion HF Buffer, 1 unit of Phusion DNA polymerase (NEB), 200 µM dNTPs, and 1 µM of each unique forward and reverse Illumina barcoding primer pair. Sequencing was performed on an Illumina MiSeq. ONE-seq sequencing data were analyzed by custom Python scripts as previously described^46, 47^.

### CRISPRme

CRISPRme^44^ (crisprme.di.univr.it) was used to perform *in silico* identification of potential off-targets by sequence similarity. We performed searches for sg1617, sg1450, sg1618 and sg1449 with no PAM restriction (NNN), up to 6 mismatches, and up to 2 DNA or RNA bulges. We used the cutting frequency determination (CFD) score^82^ to prioritize CRISPRme-nominated off-target sites and aimed to test all sites with CFD ≥ 0.4 either based on the hg38 reference genome or based on alternative alleles from variants in the 1000 Genomes Project database^83^ for which we had informative donors available. CFD score is a metric of predicted cleavage potential (ranging from 0 to 1 where the on-target has a score of 1) that was empirically determined for SpCas9 and has been shown to perform well in prioritizing off-targets that could be validated by targeted sequencing, with higher scores indicating greater likelihood of cleavage^84^. Additionally, the CFD score is suitable for sites with DNA or RNA bulges in addition to mismatches^84^.

### rhAmpSeq

rhAmpSeq assays were designed and synthesized by IDT. Off-target sites were amplified with rhAmpSeq Library Mix 1 (IDT) and using rhAmpSeq forward and reverse assay primer pools. The cycling conditions were: 95◌֯ C for 10 min; 14 cycles of 95◌֯ C for 15 s and 61◌֯ C for 8 min, and 99.5◌֯ C for 15 min. Locus specific PCR product was diluted to 1:20 and 11 µl was used for the indexing PCR with the cycling conditions: 95◌֯ C for 3 min; 24 cycles of 95◌֯ C for 15 s, 60◌֯ C for 30 s and 72◌֯ C for 30 s; and 72◌֯ C for 1 min. The resulting PCR products were evaluated by Qubit dsDNA HS Assay Kit (Thermo Fisher, Q32854), TapeStation with High Sensitivity D1000 Reagents (Agilent, 5067-5585) and High Sensitivity D1000 ScreenTape (Agilent, 5067-5584) and KAPA Universal qPCR Master Mix (KAPA Biosystems, KK4824/Roche 07960140001). The products were pooled as equimolar and subjected to deep sequencing using MiniSeq (Illumina). Amplicons were sequenced using paired-end ∼150 bp reads on Illumina HiSeq and MiniSeq instruments. Reads were trimmed for adapters and quality using Trimmomatic^75^ in paired-end mode. Editing outcomes were analyzed using CRISPResso2^76^ using the default (Cas9) parameters along with the ignore_substitutions option. CRISPRessoPooled was used for analysis of all candidate off-targets. For sites that did not achieve adequate coverage, we attempted targeted amplicon sequencing.

### Sequencing data analysis

This analysis was designed to detect and quantify off-target editing activity at a set of 1078 genomic loci nominated as candidate off-target sites for gRNAs sg1617, sg1450, sg1449 and sg1618 as described above. This nomination included candidate off-target sites from the reference genome as well as off-target sites nominated with alternative alleles if donors were available. CD34+ HSPCs from 8 healthy donors (5 donors edited with 3xNLS- SpCas9:sg1617+sg1450 and 3 donors edited with 3xNLS-HiFi-SpCas9:sg1617+sg1450) were used for sg1617 and sg1450 off-target analysis; CD34+ HSPCs from 3 healthy donors edited with 3xNLS-SpCas9:sg1618+sg1450 were used for sg1618 off-target analysis; CD34+ HSPCs from 2 healthy donors and one SCD patient donor edited with 3xNLS-HiFi-SpCas9:sg1449 were used for sg1449 off-target analysis. For each donor, some cells were untreated to provide an estimate of the sequencing background noise associated with the specific locus and donor analyzed.

Using genomic DNA extracted from these cells, we performed pooled amplicon sequencing for all candidate off-target sites via rhAmpSeq (primer designs in **Tables S5-S9**) following the standard rhAmpSeq Library Preparation Protocol (Integrated DNA Technologies). Amplicons were sequenced using paired-end ∼150 bp reads on Illumina HiSeq and MiniSeq instruments. Reads were trimmed for adapters and quality using Trimmomatic^75^ in paired-end mode unless otherwise noted in.

Editing outcomes were analyzed using CRISPResso2^76^ with default (Cas9) parameters along with the `ignore_substitutions` option. CRISPRessoPooled was used for analysis of all candidate off-targets except those for which single-end reads were used and/or were amplified individually, which were instead analyzed on a per-target basis using CRISPRessoBatch.

Single-end reads were used for loci with high background in the control samples due to nearby difficult-to-sequence homopolymers and/or read merging issues. If CRISPResso2 failed to distinguish between reference and alternative allele reads for a heterozygous donor where the expected difference is only a single nucleotide, manual filtering was performed on reads to only count those containing the SNP nucleotide. The results at loci with high sequence similarity at both the candidate off-target motif and surrounding genomic sequence were merged because the amplicons could not be easily distinguished from each other. Then, for each sample, the result at every candidate off-target site was summarized in terms of a *TotalCount*, the total number of reads aligning to that locus which satisfied quality criteria; and *IndelsCount*, the number of reads containing an indel overlapping the potential SpCas9 cleavage site (3 bp upstream of the PAM).

We modeled the proportion of *IndelsCount* over *TotalCount* using a binomial response generalized linear model (logistic regression) including the following variables as covariates: *DonorID*, a factor distinguishing the cell donor; and *Treatment,* a binary variable with value 0 for control and 1 for edited samples. The model formula is given by: Current methods for off-target site nomination are known to have high false-positive rates^84, 85^, and we intentionally set loose thresholds for off-target site nomination so we could more comprehensively assess potential genotoxicity. Hence, we anticipated observing *IndelsCount* values equal to or close to 0 in many cases. When the probability of success is very low and/or the input data table is sparse (*IndelsCount=0* for multiple samples), the estimation of logistic regression parameters can be problematic, and standard techniques are prone to fail. To address this issue, we adopted the bias-reduction approach developed by Firth^86^, a second-order unbiased estimator that returns finite standard errors for the coefficients under all conditions.

We performed the regression using R statistical software and the libraries *brglm2*^87^ and *ggeffects*. We analyzed the results for each locus separately, based on two steps:

1. Identifying validated off-targets: We considered a candidate off-target as a validated true positive if the *Treatment* coefficient is positive and significant (Wald test for the null hypothesis = 0 and α = 0.05) and the estimated *Indel* proportion in the edited sample is at least 0.1% greater than that in the paired control sample in the mean of informative donors. We corrected the p-values for multiple hypothesis testing using the FDR method^88^.
2. Determining sufficient sequencing depth for each sample: We calculated each sample’s estimated proportion of insertions and deletions (Indels) and their 95% Confidence Intervals (CI). For non-significant off-targets, we considered the sequencing depth sufficient if the difference between the CI upper estimate of Indels for the edited sample and the CI lower estimate for the control sample was less than 0.2% (provided that the edited sample had more Indels than the control sample). This sequencing coverage would allow for detecting an increment in the Indels proportion of at least 0.1% in an edited sample, assuming up to 0.1% Indels in the paired control sample (which is typical for Illumina sequencing background frequency). This method allows more rigorous coverage sufficiency assessment than an arbitrary read threshold while accounting for relationships within the two treatment groups and the paired samples for each donor.

### Micronucleus analysis

HSPCs received 0, 24 or 48 hours of pre-stimulation prior to electroporation. Cells were cultured for 48 hours after electroporation before micronucleus analysis. MicroFlow In Vitro-250/50 Kit (Litron, cat#1131099) was used for micronucleus analysis. 500,000 cells were stained with 300 μl of Nucleic Acid Dye A working solution and incubated on ice under cool white fluorescent light for 30 min. After washing, the cells were incubated in 500 μl of Complete Lysis vSolution 1 for one hour at room temperature protected from light. 500 μl of Complete Lysis Solution 2 was added and incubated for 30 min at room temperature, protected from light. Flow cytometric analysis was performed by BD FACSLyric Flow Cytometry System. The measurements excluded debris and other spurious events identified by anomalous light scatter profile and/or EMA-positive staining characteristic. Micronuclei events were defined by events possessing 1/100th to 1/10th the SYTOX Green fluorescent intensity of 2n nuclei. The fraction of micronuclei was expressed as percent by dividing the number of events that fell within the micronuclei gate by the sum of events within the nuclei plus micronuclei gates and multiplying by 100. Nuclei and micronuclei were sorted by BD FACSMelody. Enrichment of chr2p segment telomeric to *BCL11A* cleavage site was determined by ddPCR *ROCK2* assay (BioRad, assay ID: dHsaCP2500273) and reference chr1 *AGO1* assay (BioRad, assay ID: dHsaCP2500349).

After droplet generation, PCR followed the cycling conditions: 1 cycle of 95°C for 10 min, 50 cycles of 94°C for 30 sec, 58°C for 60 sec, 1 cycle of 98°C for 10 min, held at 4°C. PCR product was read by droplet reader and the results were analyzed using the QuantaSoft software.

### In vitro sickling assay

*In vitro* differentiated erythroid cells were harvested at the terminal differentiation on day 18 and stained with 2 μg/ml of Hoechst 33342 (Millipore, cat# B2261) for 15 min at 37°C in the dark.

Hoechst negative enucleated cells were sorted by BD FACSAria II. 500,000 enucleated cells in 500 μl of EDM medium were treated with 500 μl of 1.5% of sodium metabisulfite (MBS) (Sigma, cat #08982-1G) in a 24-well plate for 30 min at room temperature. Live cell images were acquired using a Nikon Eclipse Ti inverted microscope. Fraction of sickled cells were calculated as the number of sickled cells divided by the number of total cells, counting at least 200 cells.

### Cell cycle analysis

Human CD34+ HSPCs immediately after thawing or selection, 24 hours and 48 hours after cytokine culture were resuspended in SCGM media at the concentration of 1,000,000/ml, stained with 2 μg/ml of Hoechst 33342 (Millipore, cat# B2261) for 45 min at 37°C in the dark, mixing every 15 min. Pyronin Y (Sigma, cat #83200-10G) was added to the cells to final concentration 5 μg/ml and incubated for 45 min at 37°C in the dark, mixing every 15 min. Cell cycle was analyzed by BD FACSLyric Flow Cytometry System and data were analyzed with FlowJo™ software.

## SUPPLEMENTAL INFORMATION

Supplemental information includes seven figures and nine tables.

## ACKNOWLEDGMENTS

The authors thank Yuxuan Wu, Erica B. Esrick, Hye Ji Cha, Stuart H. Orkin, Jasmine Bonanno, Johnathan Tran, Anne H. Shen, Jonathan Y. Hsu, Samuele Cancellieri, Manuel Tognon, Rosalba Giugno, Tara Ellison, Olivier Negre, Gabor Veres, Lauryn Christiansen and the HSCI-BCH Flow Cytometry Research Lab for technical support and helpful discussions. CD34+ HSPCs were provided by Fred Hutch Cooperative Center of Excellence in Hematology (U54 DK106829). K.P. was funded by the Deutsche Forschungsgemeinschaft (DFG, German Research Foundation), Projektnummer 417577129. L.P. was supported in part by the National Institutes of Health (NIH) (R35 HG010717). J.K.J was supported by the Defense Advanced Research Projects Agency (HR0011-17-2-0042), the U.S. National Institutes of Health (NIH) (RM1 HG009490 and R35 GM118158), the Desmond and Ann Heathwood MGH Research Scholar Award, and the Robert B. Colvin, M.D. Endowed Chair in Pathology. S.A.W. was supported in part by the National Institutes of Health (R01HL120669, OT2HL154984) and Doris Duke Charitable Foundation. D.E.B. was supported by the National Heart, Lung, and Blood Institute (OT2HL154984, P01HL053749), Burroughs Wellcome Fund, Harvard Stem Cell Institute, Doris Duke Charitable Foundation and the St. Jude Children’s Research Hospital Collaborative Research Consortium.

## AUTHOR CONTRIBUTIONS

Conceptualization, J.Z., S.D., J.F.T., S.A.W., D.E.B.; Methodology, D.E.B., S.A.W., M.A., C.B., D.A.W., J.P.M., L.P., J.K.J. V.P. and S.M.P.; Investigation, J.Z., M.N., M.F.C., N.R.N., D.A., E.M., S.A.M., T.R. D.G.J. and S.N.P.; Formal analysis, A.V., L.Y.L., L.F., P.L., D.P., L.J.Z., K.P., V.P. and K.C.; Writing – Original Draft, D.E.B. and J.Z.; Writing – Review & Editing, D.E.B., J.Z. and all authors; Funding Acquisition, D.E.B; Resources, D.E.B., M.A., J.K.J. and S.A.W.; Supervision, D.E.B.

---

**Figure S1.**
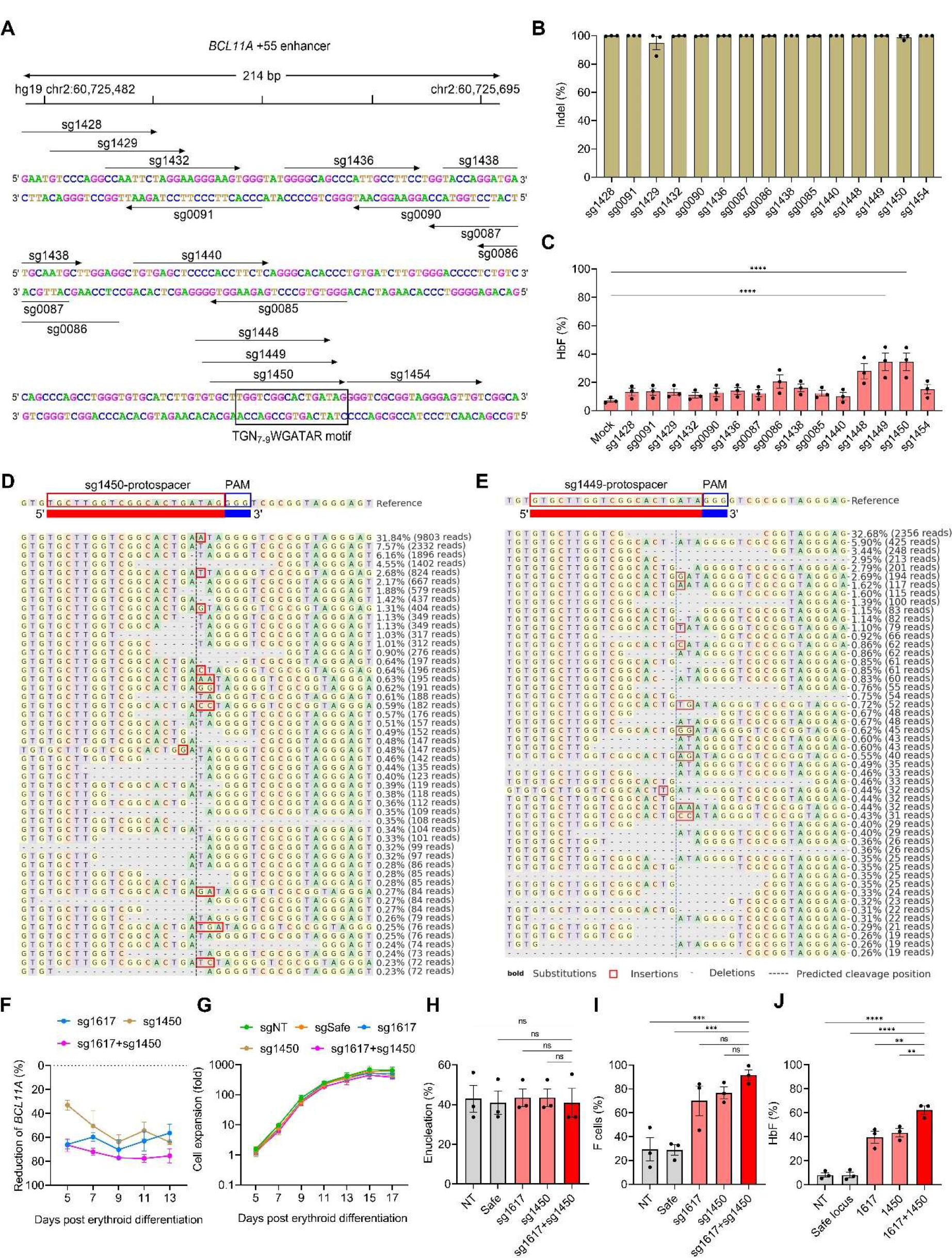
Efficient editing and HbF induction by targeting the +55 *BCL11A* erythroid enhancer in CD34+ HSPCs (A) Fifteen sgRNAs were designed to target core +55 *BCL11A* enhancer. Half E-Box/GATA (TGN7-9WGATAR) motifs which are binding sites for erythroid transcription factors TAL1 and GATA1 marked in the frame. (B) Indel frequency of 3xNLS-SpCas9 complexed with various gRNAs in CD34+ HSPCs measured by TIDE analysis. Data are plotted as mean ± SEM, n = 3 healthy donors (HD 4, HD 5 and HD6). (C) HbF induction by HPLC analysis in erythroid cells *in vitro* differentiated from RNP edited CD34+ HSPCs from three healthy donors (HD 4, HD 5 and HD 6). Data are plotted as mean ± SEM and analyzed with the unpaired two-tailed Student’s *t*-test. ns: nonsignificant, \*\*\*\**P* < 0.0001. n = 3 healthy donors. (D) Indels analyzed by CRISPResso 2 following 3xNLS-Cas9: sg1449 RNP electroporation of CD34+ HSPCs. (E) Indels analyzed by CRISPResso 2 following 3xNLS-Cas9: sg1450 RNP electroporation of CD34+ HSPCs. (F) Reduction of *BCL11A* mRNA expression compared to safe locus editing by ddPCR in erythroid precursors. Data are plotted as mean ± SEM, n = 3 healthy donors (HD 1, HD 2 and HD 3). (G) Cell expansion during erythroid differentiation. Data are plotted as mean ± SEM, n = 3 healthy donors (HD 1, HD 2 and HD 3). (H) Enucleation by Hoechst staining at the terminal erythroid differentiation. Data are plotted as mean ± SEM, n = 3 healthy donors (HD 1, HD 2 and HD 3). (I) Fraction of F cells by flow cytometry analysis. Data are plotted as mean ± SEM and analyzed with one-way ANOVA. ns: nonsignificant. n = 3 healthy donors (HD 1, HD 2 and HD 3). (J) HbF levels by HPLC in *in vitro* differentiated erythroid cells from three healthy donors (HD 1, HD 2 and HD 3). Data are plotted as mean ± SEM and analyzed with one-way ANOVA. ns: nonsignificant, \*\*\**P* < 0.001.

**Figure S2.**
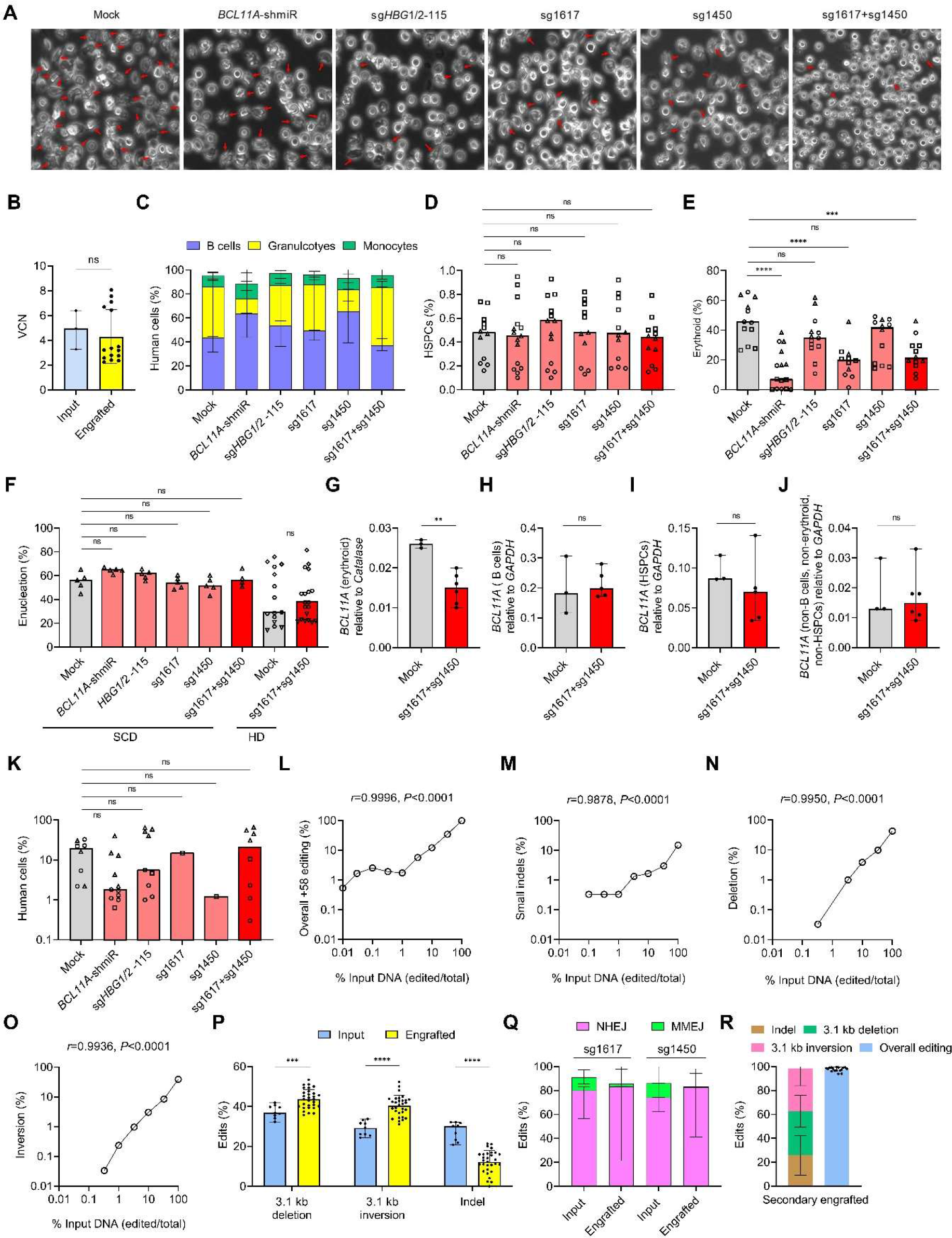
Combined editing of +58 and +55 *BCL11A* erythroid enhancers was compatible with hematopoietic repopulation in primary and secondary xenotransplant (A) Phase-contrast microscope imaging of enucleated *in vitro* differentiated erythroid cells from SCD donor after MBS treatment. (B) VCN by qPCR after lentiviral erythroid-restricted shRNA targeting *BCL11A* transduction in input cells and engrafted cells. Data are plotted as median with range and analyzed with the unpaired two-tailed Student’s *t*-test. ns: nonsignificant. n = 3 (HD 7, SCD 1 and SCD 2) for input, n = 15 primary recipients for engrafted. (C) Multilineage repopulation in BM 16 weeks after transplantation. Data are plotted as median with range. n = 12-15 primary recipients infused with HSPCs from HD 7, SCD 1 and SCD 2. (D) HSPCs engraftment in BM 16 weeks after transplantation. Data are plotted as grand median and analyzed with one-way ANOVA. ns: nonsignificant. n = 12-15 primary recipients infused with HSPCs from HD 7, SCD 1 and SCD 2. (E) Erythroid engraftment in BM 16 weeks after transplantation. Data are plotted as grand median and analyzed with one-way ANOVA. ns: nonsignificant. \*\*\**P* < 0.001, \*\*\*\**P* < 0.0001. n = 12-15 primary recipients infused with HSPCs from HD 7, SCD 1 and SCD 2. (F) Enucleation by flow analysis of Hoechst staining in engrafted erythroid from recipients infused with human HSPCs from four healthy donors and one SCD patient (HD 9, HD 10, HD 11 and SCD 2). Data are plotted as grand median and analyzed with one-way ANOVA. ns: nonsignificant. n = 4-12 primary recipients. (G-J) *BCL11A* expression by RT-qPCR in engrafted bone marrow human erythroid (G), B cell (H), HSPC (I) and non-B cells, non-erythroid, non-HSPCs (J). Data are plotted as median with range and analyzed with unpaired two-tailed Student’s *t*-test. ns: nonsignificant, \*\**P* < 0.01. n = 3-5 primary recipients infused with HSPCs from HD 11. (K) Human chimerism 16 weeks after secondary transplantation. Data are plotted as grand median and analyzed with one-way ANOVA. ns: nonsignificant. n = 1-11 secondary recipients infused BM from the primary recipients from HD 7, SCD 1 and SCD 2. (L-O) Overall +58 edit (L), small indels (M), deletion (N) and inversion (O) of edited input DNA mixed with unedited DNA in different ratios measured by +58 drop-off ddPCR, +58 offset ddPCR, 3.1 kb deletion ddPCR and 3.1 kb inversion ddPCR assays. Data are analyzed with the two-tailed nonparametric Spearman correlation. n = 3 technical replicates. (P) Frequencies of deletion, inversion and indel by ddPCR assays in engrafted cells compared to input cells following sg1617+sg1450 editing from 8 healthy donors and 2 SCD donors (HD 7, HD 9-HD 15, SCD 1 and SCD 2) . Data are plotted as median with range and analyzed with the unpaired two-tailed Student’s *t*-test. \*\**P* < 0.01, \*\*\*\**P* < 0.0001. (Q) Edited alleles repaired by NHEJ (-8 to +6bp) and MMEJ (-9 to -20bp) in engrafted BM cells as compared to input cells, analyzed by Sanger sequencing. Data are plotted as median with range. n = 10 donors (HD 7, HD 9-HD 15, SCD 1 and SCD 2) for input, n = 41-42 primary recipients for engrafted. (R) Frequencies of deletion, inversion, indel and overall edits by ddPR assays in secondary engrafting cells following sg1617+sg1450 editing. Data are plotted as median with range. n = 17 secondary recipients infused with primary BM from SCD 1, SCD 2, HD 7, HD 9, HD 10 and HD 13.

**Figure S3.**
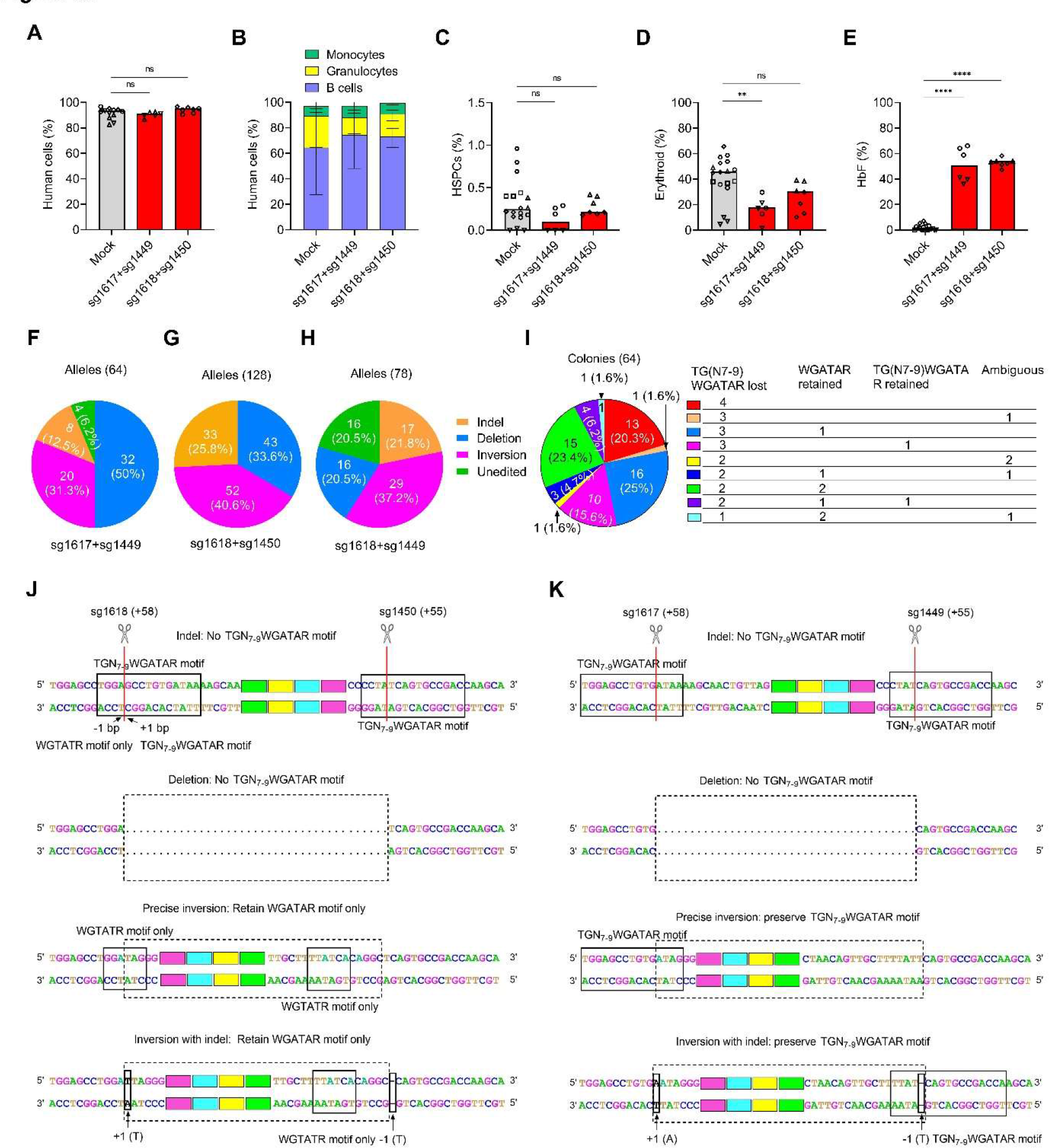
Alternate pairs of combined +58 and +55 editing achieve similar editing efficiency, motif disruption, and HbF induction (A) Human chimerism in BM 16 weeks after infusion with human CD34+ HSPCs edited with sg1617+sg1449 and sg1618+sg1450. Data are plotted as grand median and analyzed with the unpaired two-tailed Student’s *t*-test. ns: nonsignificant. n = 6-18 primary recipients infused with HSPCs from HD 16 and HD 17 for sg1617+1449, and HD 7 and HD 13 for sg1618+sg1450. (B) Multilineage repopulation in BM 16 weeks after infusion with human CD34+ HSPCs edited with sg1617+sg1449 from HD 16 and HD 17 and sg1618+sg1450 from HD 7 and HD 13. Data are plotted as median with range. n = 6-18 primary recipients. (C and D) HSPCs (C) and erythroid (D) engraftment in BM 16 weeks after infusion with human CD34+ HSPCs edited with sg1617+sg1449 from HD 16 and HD 17 and sg1618+sg1450 from HD 7 and HD 13. Data are plotted as grand median and analyzed with the unpaired two-tailed Student’s *t*-test. ns: nonsignificant, \*\**P* < 0.01. (E) HbF levels by HPLC in engrafted BM erythroid cells edited with sg1617+sg1449 from HD 16 and HD 17 and sg1618+sg1450 from HD 7 and HD 13. Data are plotted as grand median and analyzed with the unpaired two-tailed Student’s *t*-test. \*\*\*\**P* < 0.0001. (F-H) Allelic analysis of single clones after sg1617+sg1449 editing (F) derived from HD 16 and HD 17, sg1618+sg1450 editing (G) from HD 8 and sg1618+sg1449 editing (H) from HD 28. (I) Junctional analysis of TGN7-9WGATAR half E-box/GATA binding motifs after sg1618+sg1450 editing in individual single cell derived erythroid colonies after editing of CD34+ HSPCs from HD 8. (J and K) Schema of edit outcomes after sg1618+sg1450 editing (J), sg1617+sg1449 editing (K) and their associated disruption/preservation of TGN7-9WGATAR half E-box/GATA binding motifs.

**Figure S4.**
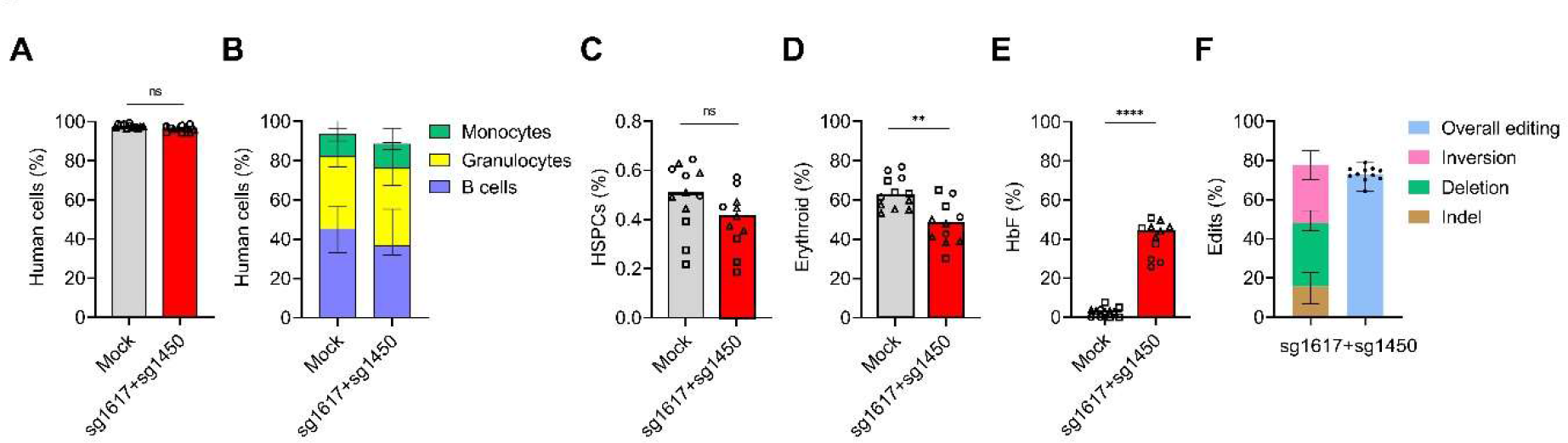
Robust engraftment and HbF induction following 3xNLS-HiFi- SpCas9:sg1617+sg1450 gene editing of CD34+ HSPCs. (A) Human chimerism 16 weeks after infusion with 3xNLS-HiFi-SpCas9:sg1617+sg1450 edited HSPCs from 3 healthy donors (HD19, HD 20 and HD 21). Data are plotted as grand median and analyzed with the unpaired two-tailed Student’s *t*-test. ns: nonsignificant. n = 11-12 primary recipients. (B) The engraftment of B cell, granulocyte, monocyte in bone marrow 16 weeks after infusion with 3xNLS-HiFi-SpCas9:sg1617+sg1450 edited HSPCs from 3 healthy donors (HD19, HD 20 and HD 21). Data are plotted as median with range. n = 11-12 primary recipients. (C and D) The engraftment of HSPCs (C) or erythroid cells (D) in the bone marrow 16 weeks after infusion with 3xNLS-HiFi-SpCas9:sg1617+sg1450 edited HSPCs from 3 healthy donors (HD19, HD 20 and HD 21). Data are plotted as grand median and analyzed with the unpaired two-tailed Student’s *t*-test. ns: nonsignificant, \*\**P* < 0.01. n = 11-12 primary recipients. (E) HbF levels by HPLC analysis from engrafted erythroid cells. Data are plotted as grand median and analyzed with the unpaired two-tailed Student’s *t*-test. \*\*\*\**P* < 0.0001. n = 11-12 primary recipients infused with edited HSPCs from HD 19, HD 20 and HD 21. (F) Frequency of 3 kb deletion, inversion, indel and overall edits by ddPCR in engrafted cells infused with edited HSPCs from HD19, HD 20 and HD 21. Data are plotted as median with range. n = 11 primary recipients.

**Figure S5.**
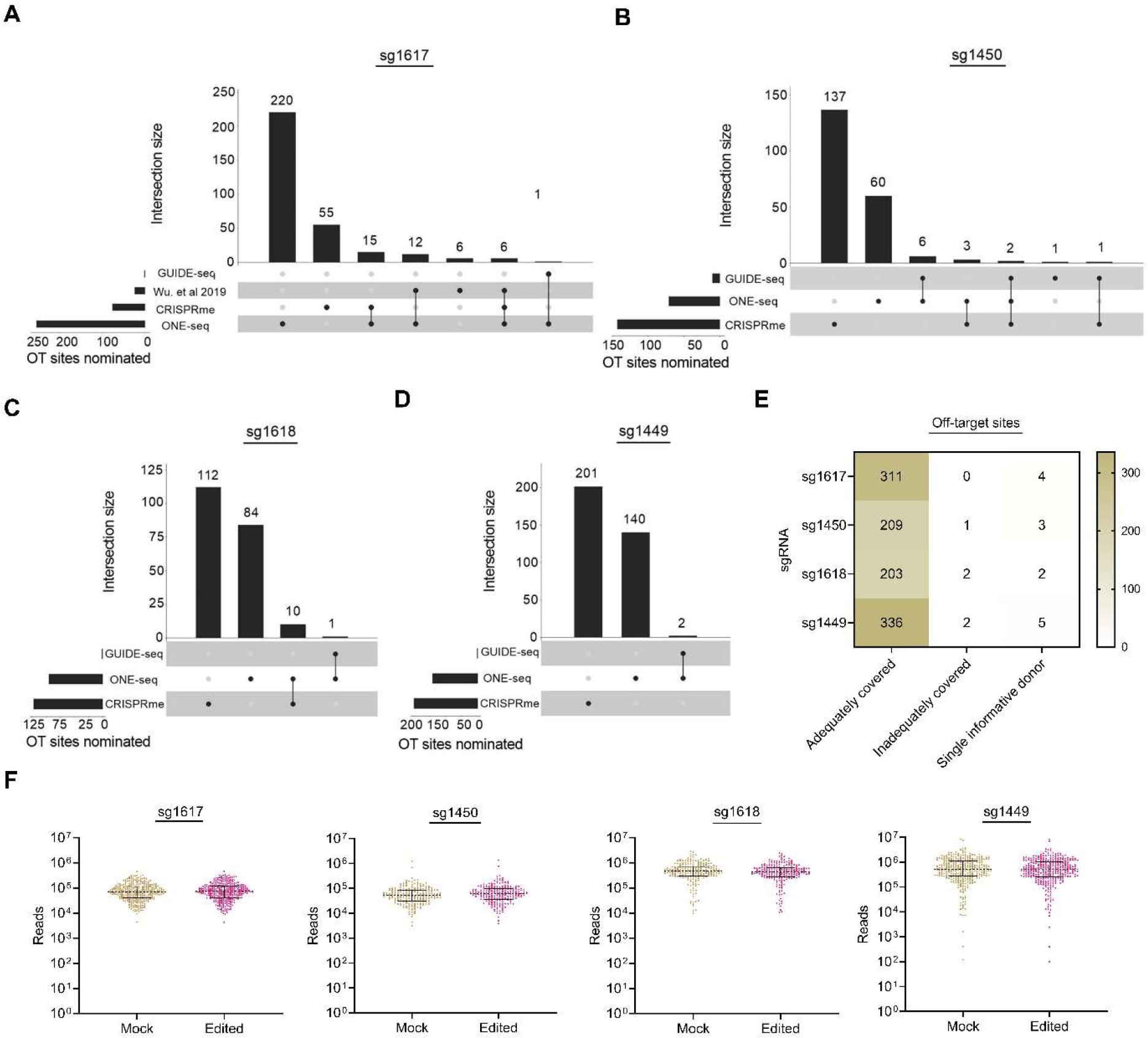
Nomination of candidate off-target sites (A-D) Candidate off-target sites nominated by *in silico* prediction with Cas-OFFinder, GUIDE- seq, CRISPRme, and ONE-seq for sg1617 (A), sg1450 (B), sg1618 (C), and sg1449 (D). GUIDE-seq was performed in HSPC donors, HD 22, HD 23 and HD 24 for sg1617 and sg1450; HD 25, HD 26 and HD 27 for sg1618; HD 3, HD 28 and HD 29 for sg1449. (E) Number of candidate off-target sites based on three categories: adequately covered with the difference between CI upper estimate of indels for the edited sample and the CI lower estimate for the control sample (delta-CI) <0.2%, inadequately covered (delta-CI≥0.2%), and, for variant- associated off-target sites, sites with a single informative donor for which delta-CI could not be calculated. (F) Sequencing reads per candidate off-target site for sg1617, sg1450, sg1618 and sg1449. Each dot represents a unique off-target site. Lines represent median and interquartile range.

**Figure S6.**
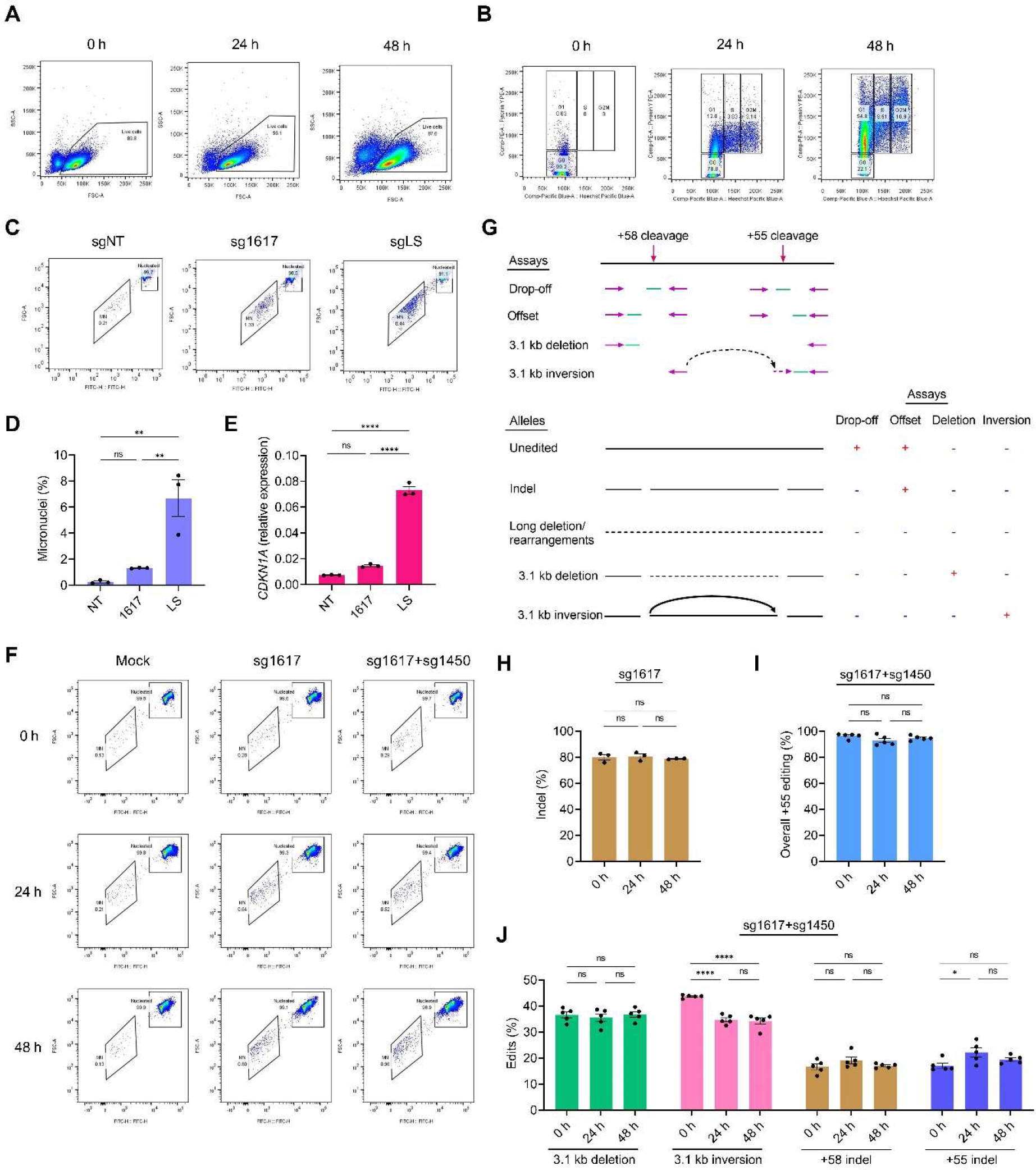
Measurements of micronucleation and long deletion/rearrangements (A) Cell size at 0h, 24 h and 48 h post pre-stimulation by forward scatter. (B) Cell cycle analysis by flow after Pyronin Y and Hoechst staining. (C) Micronucleus analysis by flow for CD34+ HSPCs electroporated with Cas9:sgNT (non- targeting) RNP, Cas9:sg1617 RNP, Cas9:sgLS (low specificity) RNP after 48 hours in cytokine culture. (D) Frequency of micronucleation 48 hours following gene editing. Data are plotted as mean ± SEM and analyzed with one-way ANOVA, ns: nonsignificant, \*\**P* < 0.01. n = 3 healthy donors (HD 26, HD 27 and HD 30). (E) RT-qPCR analysis of *p21* expression relative to *GAPDH* 5 h after gene editing. Data are plotted as mean ± SEM and analyzed with one-way ANOVA, ns: nonsignificant, \*\*\*\**P* < 0.0001. n = 3 healthy donors (HD 26, HD 27 and HD 30). (F) Micronucleus analysis by flow for CD34+ HSPCs electroporated with SpCas9:sg1617 and SpCas9:sg1617+sg1450 RNP at 0, 24, or 48 hours in cytokine culture. (G) The schema of ddPCR assays. +58 drop-off ddPCR assay measures the overall +58 editing including 3.1kb deletion, 3.1kb inversion, +58 indels and long deletion/rearrangements. +55 drop-off ddPCR assay measures the overall +55 editing including 3.1kb deletion, 3.1kb inversion, +55 indels and long deletion/rearrangements. (H) Frequency of indel by +58 drop-off and +58 offset ddPCRs following sg1617 single editing. Data are plotted as mean ± SEM and analyzed with one-way ANOVA, ns: nonsignificant. n = 3 healthy donors (HD 32, HD 33 and HD 34). (I) Overall +55 editing by +55 drop-off ddPCR assay following sg1617+sg1450 editing. Data are plotted as mean ± SEM and analyzed with one-way ANOVA, ns: nonsignificant. n = 5 healthy donors (HD 2, HD 3 and HD 32-HD 34). (J) Frequency of deletion, inversion and +58 indel and +55 indel after sg1617+sg1450 editing by ddPCR assays. Data are plotted as mean ± SEM and analyzed with one-way ANOVA, ns: nonsignificant, \**P* < 0.05, \*\*\*\**P* < 0.0001. n = 5 healthy donors (HD 2, HD 3 and HD 32-HD 34).

**Figure S7.**
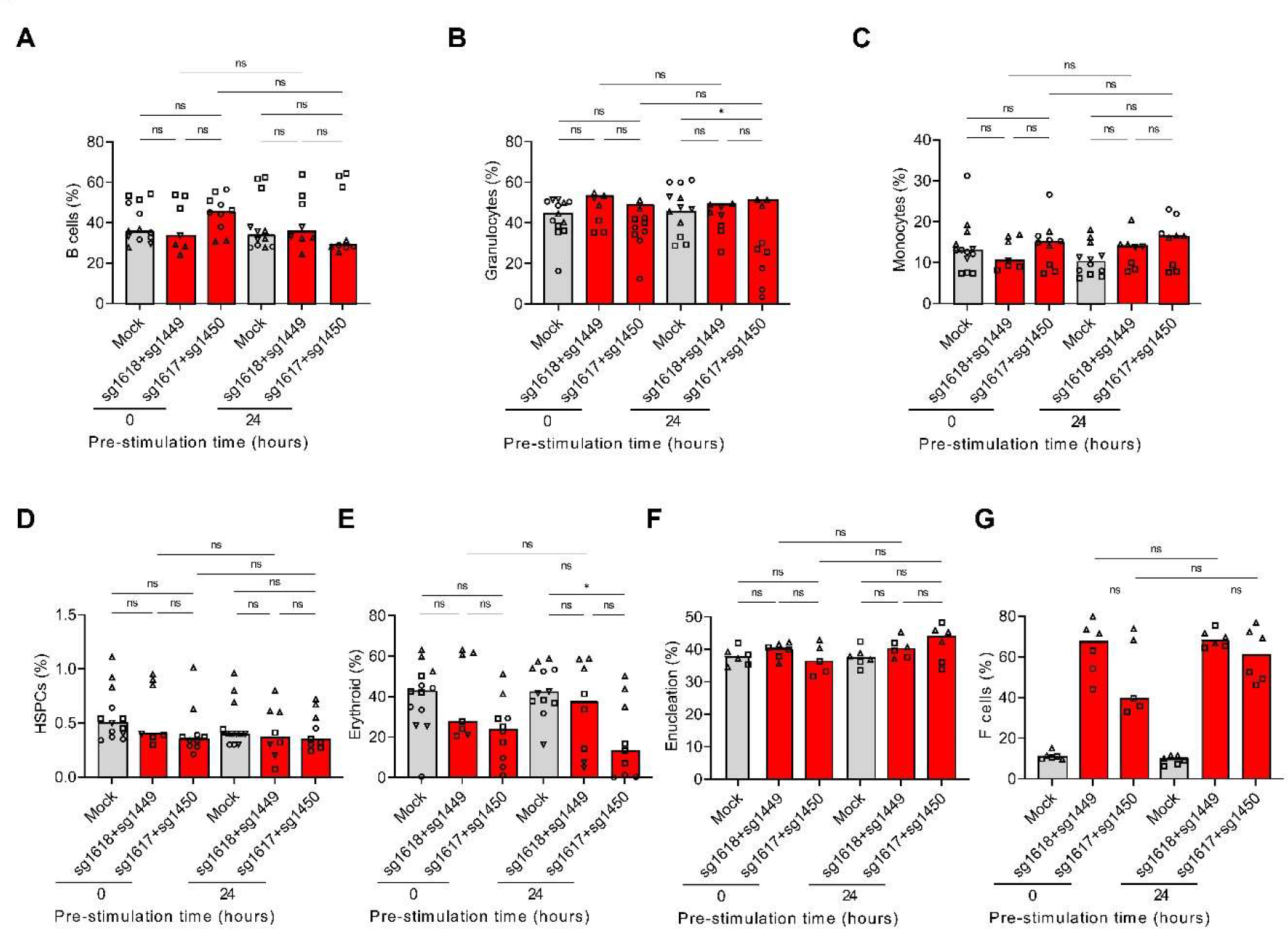
Intact hematopoietic repopulation by HSPCs edited without *ex vivo* cytokine culture. (A-E) B cell (A), granulocyte (B), monocyte (C), HSPC (D) and erythroid (E) engraftment 16 weeks after transplantation. Data are plotted as grand median and analyzed with one-way ANOVA, ns: nonsignificant, \**P* < 0.5, n = 7-13 primary recipients infused with HSPCs from HD 28, HD29, HD 35 and SCD 3. (F) Enucleation of engrafted erythroid cells isolated by human CD235a microbeads. Data are plotted as grand median and analyzed with one-way ANOVA, ns: nonsignificant. n = 5-6 primary recipients infused with HSPCs from HD 29 and SCD 3. (G) Fraction of F cells in engrafted enucleated erythroid cells. Data are plotted as grand median and analyzed with one-way ANOVA, ns: nonsignificant. n = 5-6 primary recipients infused with HSPCs from HD 29 and SCD 3.

## REFERENCES

1. Piel, F.B., Steinberg, M.H., and Rees, D.C. (2017). Sickle Cell Disease. N. Engl. J. Med. 377, 305.

2. Motta, I., Ghiaccio, V., Cosentino, A., and Breda, L. (2019). Curing Hemoglobinopathies: Challenges and Advances of Conventional and New Gene Therapy Approaches. Mediterr. J. Hematol. Infect. Dis. 11, e2019067.

3. Orkin, S.H., and Bauer, D.E. (2019). Emerging Genetic Therapy for Sickle Cell Disease. Annu. Rev. Med. 70, 257–271.

4. Christakopoulos, G.E., Telange, R., Yen, J., and Weiss, M.J. (2023). Gene Therapy and Gene Editing for β-Thalassemia. Hematol. Oncol. Clin. North Am. 37, 433–447.

5. Lettre, G., and Bauer, D.E. (2016). Fetal haemoglobin in sickle-cell disease: from genetic epidemiology to new therapeutic strategies. Lancet 387, 2554–2564.

6. Frangoul, H., Altshuler, D., Cappellini, M.D., Chen, Y.-S., Domm, J., Eustace, B.K., Foell, J., de la Fuente, J., Grupp, S., Handgretinger, R., et al. (2021). CRISPR-Cas9 Gene Editing for Sickle Cell Disease and β-Thalassemia. N. Engl. J. Med. 384, 252–260.

7. Esrick, E.B., Lehmann, L.E., Biffi, A., Achebe, M., Brendel, C., Ciuculescu, M.F., Daley, H., MacKinnon, B., Morris, E., Federico, A., et al. (2021). Post-Transcriptional Genetic Silencing of BCL11A to Treat Sickle Cell Disease. N. Engl. J. Med. 384, 205–215.

8. Sales, R.R., Nogueira, B.L., Tosatti, J.A.G., Gomes, K.B., and Luizon, M.R. (2021). Do Genetic Polymorphisms Affect Fetal Hemoglobin (HbF) Levels in Patients With Sickle Cell Anemia Treated With Hydroxyurea? A Systematic Review and Pathway Analysis. Front. Pharmacol. 12, 779497.

9. Belisário, A.R., Sales, R.R., Silva, C.M., Velloso-Rodrigues, C., and Viana, M.B. (2016). The Natural History of Hb S/Hereditary Persistence of Fetal Hemoglobin in 13 Children from the State of Minas Gerais, Brazil. Hemoglobin 40, 215–219.

10. Talbot, J.F., Bird, A.C., and Serjeant, G.R. (1983). Retinal changes in sickle cell/hereditary persistence of fetal haemoglobin syndrome. Br. J. Ophthalmol. 67, 777– 778.

11. Whyte, D., Forget, B., Chui, D.H.K., Luo, H.-Y., and Pashankar, F. (2013). Massive splenic infarction in an adolescent with hemoglobin S-HPFH. Pediatr. Blood Cancer 60, E49–E51.

12. Edington, G.M., and Lehmann, H. (1955). Expression of the sickle-cell gene in Africa. Br. Med. J. 1, 1308–1311.

13. Jacob, G.F., and Raper, A.B. (1958). Hereditary persistence of foetal haemoglobin production, and its interaction with the sickle-cell trait. Br. J. Haematol. 4, 138–149.

14. Conley, C.L., Weatherall, D.J., Richardson, S.N., Shepard, M.K., and Charache, S. (1963). Hereditary persistence of fetal hemoglobin: a study of 79 affected persons in 15 Negro families in Baltimore. Blood 21, 261–281.

15. Maier-Redelsperger, M., Noguchi, C.T., de Montalembert, M., Rodgers, G.P., Schechter, A.N., Gourbil, A., Blanchard, D., Jais, J.P., Ducrocq, R., and Peltier, J.Y. (1994). Variation in fetal hemoglobin parameters and predicted hemoglobin S polymerization in sickle cell children in the first two years of life: Parisian Prospective Study on Sickle Cell Disease. Blood 84, 3182–3188.

16. Henry, E.R., Cellmer, T., Dunkelberger, E.B., Metaferia, B., Hofrichter, J., Li, Q., Ostrowski, D., Ghirlando, R., Louis, J.M., Moutereau, S., et al. (2020). Allosteric control of hemoglobin S fiber formation by oxygen and its relation to the pathophysiology of sickle cell disease. Proc. Natl. Acad. Sci. U. S. A. 117, 15018–15027.

17. Porteus, M.H. (2019). A New Class of Medicines through DNA Editing. N. Engl. J. Med. 380, 947–959.

18. Kieffer, S.R., and Lowndes, N.F. (2022). Immediate-Early, Early, and Late Responses to DNA Double Stranded Breaks. Front. Genet. 13, 793884.

19. Iyer, S., Suresh, S., Guo, D., Daman, K., Chen, J.C.J., Liu, P., Zieger, M., Luk, K., Roscoe, B.P., Mueller, C., et al. (2019). Precise therapeutic gene correction by a simple nuclease-induced double-stranded break. Nature 568, 561–565.

20. Genovese, P., Schiroli, G., Escobar, G., Di Tomaso, T., Firrito, C., Calabria, A., Moi, D., Mazzieri, R., Bonini, C., Holmes, M.C., et al. (2014). Targeted genome editing in human repopulating haematopoietic stem cells. Nature 510, 235–240.

21. Kosicki, M., Tomberg, K., and Bradley, A. (2018). Repair of double-strand breaks induced by CRISPR-Cas9 leads to large deletions and complex rearrangements. Nat. Biotechnol. 36, 765–771.

22. Leibowitz, M.L., Papathanasiou, S., Doerfler, P.A., Blaine, L.J., Sun, L., Yao, Y., Zhang, C.-Z., Weiss, M.J., and Pellman, D. (2021). Chromothripsis as an on-target consequence of CRISPR-Cas9 genome editing. Nat. Genet. 53, 895–905.

23. Bauer, D.E., Kamran, S.C., Lessard, S., Xu, J., Fujiwara, Y., Lin, C., Shao, Z., Canver, M.C., Smith, E.C., Pinello, L., et al. (2013). An Erythroid Enhancer of BCL11A Subject to Genetic Variation Determines Fetal Hemoglobin Level. Science 342, 253–257.

24. Canver, M.C., Smith, E.C., Sher, F., Pinello, L., Sanjana, N.E., Shalem, O., Chen, D.D., Schupp, P.G., Vinjamur, D.S., Garcia, S.P., et al. (2015). BCL11A enhancer dissection by Cas9-mediated in situ saturating mutagenesis. Nature 527, 192–197.

25. Vierstra, J., Reik, A., Chang, K.-H.H., Stehling-Sun, S., Zhou, Y., Hinkley, S.J., Paschon, D.E., Zhang, L., Psatha, N., Bendana, Y.R., et al. (2015). Functional footprinting of regulatory DNA. Nat. Methods 12, 927–930.

26. Huang, P., Peslak, S.A., Lan, X., Khandros, E., Yano, J.A., Sharma, M., Keller, C.A., Giardine, B., Qin, K., Abdulmalik, O., et al. (2020). The HRI-regulated transcription factor ATF4 activates BCL11A transcription to silence fetal hemoglobin expression. Blood 135, 2121–2132.

27. Huang, P., Peslak, S.A., Ren, R., Khandros, E., Qin, K., Keller, C.A., Giardine, B., Bell, H.W., Lan, X., Sharma, M., et al. (2022). HIC2 controls developmental hemoglobin switching by repressing BCL11A transcription. Nat. Genet. 54, 1417–1426.

28. Wu, Y., Zeng, J., Roscoe, B.P., Liu, P., Yao, Q., Lazzarotto, C.R., Clement, K., Cole, M.A., Luk, K., Baricordi, C., et al. (2019). Highly efficient therapeutic gene editing of human hematopoietic stem cells. Nat. Med. 25, 776–783.

29. Demirci, S., Zeng, J., Wu, Y., Uchida, N., Shen, A.H., Pellin, D., Gamer, J., Yapundich, M., Drysdale, C., Bonanno, J., et al. (2020). BCL11A enhancer–edited hematopoietic stem cells persist in rhesus monkeys without toxicity. J. Clin. Invest. 130, 6677–6687.

30. Fu, B., Liao, J., Chen, S., Li, W., Wang, Q., Hu, J., Yang, F., Hsiao, S., Jiang, Y., Wang, L., et al. (2022). CRISPR-Cas9-mediated gene editing of the BCL11A enhancer for pediatric β0/β0 transfusion-dependent β-thalassemia. Nat. Med. 28, 1573–1580.

31. Hsu, J.Y., Fulco, C.P., Cole, M.A., Canver, M.C., Pellin, D., Sher, F., Farouni, R., Clement, K., Guo, J.A., Biasco, L., et al. (2018). CRISPR-SURF: discovering regulatory elements by deconvolution of CRISPR tiling screen data. Nat. Methods 15, 992–993.

32. Liu, N., Xu, S., Yao, Q., Zhu, Q., Kai, Y., Hsu, J.Y., Sakon, P., Pinello, L., Yuan, G.-C., Bauer, D.E., et al. (2021). Transcription factor competition at the γ-globin promoters controls hemoglobin switching. Nat. Genet. 53, 511–520.

33. Wadman, I.A., Osada, H., Grütz, G.G., Agulnick, A.D., Westphal, H., Forster, A., and Rabbitts, T.H. (1997). The LIM-only protein Lmo2 is a bridging molecule assembling an erythroid, DNA-binding complex which includes the TAL1, E47, GATA-1 and Ldb1/NLI proteins. EMBO J. 16, 3145–3157.

34. Han, G.C., Vinayachandran, V., Bataille, A.R., Park, B., Chan-Salis, K.Y., Keller, C.A., Long, M., Mahony, S., Hardison, R.C., and Pugh, B.F. (2016). Genome-Wide Organization of GATA1 and TAL1 Determined at High Resolution. Mol. Cell. Biol. 36, 157–172.

35. Brendel, C., Guda, S., Renella, R., Bauer, D.E., Canver, M.C., Kim, Y.-J., Heeney, M.M., Klatt, D., Fogel, J., Milsom, M.D., et al. (2016). Lineage-specific BCL11A knockdown circumvents toxicities and reverses sickle phenotype. J. Clin. Invest. 126, 3868–3878.

36. Traxler, E.A., Yao, Y., Wang, Y.-D., Woodard, K.J., Kurita, R., Nakamura, Y., Hughes, J.R., Hardison, R.C., Blobel, G.A., Li, C., et al. (2016). A genome-editing strategy to treat β-hemoglobinopathies that recapitulates a mutation associated with a benign genetic condition. Nat. Med. 22, 987–990.

37. Métais, J.-Y., Doerfler, P.A., Mayuranathan, T., Bauer, D.E., Fowler, S.C., Hsieh, M.M., Katta, V., Keriwala, S., Lazzarotto, C.R., Luk, K., et al. (2019). Genome editing of HBG1 and HBG2 to induce fetal hemoglobin. Blood Adv 3, 3379–3392.

38. Liu, N., Hargreaves, V.V., Zhu, Q., Kurland, J.V., Hong, J., Kim, W., Sher, F., Macias-Trevino, C., Rogers, J.M., Kurita, R., et al. (2018). Direct Promoter Repression by BCL11A Controls the Fetal to Adult Hemoglobin Switch. Cell 173, 430–442.e17.

39. Martyn, G.E., Wienert, B., Yang, L., Shah, M., Norton, L.J., Burdach, J., Kurita, R., Nakamura, Y., Pearson, R.C.M., Funnell, A.P.W., et al. (2018). Natural regulatory mutations elevate the fetal globin gene via disruption of BCL11A or ZBTB7A binding. Nat. Genet. 50, 498–503.

40. McIntosh, B.E., Brown, M.E., Duffin, B.M., Maufort, J.P., Vereide, D.T., Slukvin, I.I., and Thomson, J.A. (2015). Nonirradiated NOD,B6.SCID Il2rγ-/- Kit(W41/W41) (NBSGW) mice support multilineage engraftment of human hematopoietic cells. Stem Cell Reports 4, 171–180.

41. Lee, H.J., Kim, E., and Kim, J.-S. (2010). Targeted chromosomal deletions in human cells using zinc finger nucleases. Genome Res. 20, 81–89.

42. Gupta, A., Hall, V.L., Kok, F.O., Shin, M., McNulty, J.C., Lawson, N.D., and Wolfe, S.A. (2013). Targeted chromosomal deletions and inversions in zebrafish. Genome Res. 23, 1008–1017.

43. Canver, M.C., Bauer, D.E., Dass, A., Yien, Y.Y., Chung, J., Masuda, T., Maeda, T., Paw, B.H., and Orkin, S.H. (2014). Characterization of Genomic Deletion Efficiency Mediated by CRISPR/Cas9 in Mammalian Cells. J. Biol. Chem. 289, 21312–21324.

44. Cancellieri, S., Zeng, J., Lin, L.Y., Tognon, M., Nguyen, M.A., Lin, J., Bombieri, N., Maitland, S.A., Ciuculescu, M.-F., Katta, V., et al. (2023). Human genetic diversity alters off-target outcomes of therapeutic gene editing. Nat. Genet. 55, 34–43.

45. Vakulskas, C.A., Dever, D.P., Rettig, G.R., Turk, R., Jacobi, A.M., Collingwood, M.A., Bode, N.M., McNeill, M.S., Yan, S., Camarena, J., et al. (2018). A high-fidelity Cas9 mutant delivered as a ribonucleoprotein complex enables efficient gene editing in human hematopoietic stem and progenitor cells. Nat. Med. 24, 1216–1224.

46. 46. Petri, K. et al (2021). Global-scale CRISPR gene editor specificity profiling by ONE-seq identifies population-specific, variant off-target effects. bioRxiv. 10.1101/2021.04.05.438458.

47. Musunuru, K., Chadwick, A.C., Mizoguchi, T., Garcia, S.P., DeNizio, J.E., Reiss, C.W., Wang, K., Iyer, S., Dutta, C., Clendaniel, V., et al. (2021). In vivo CRISPR base editing of PCSK9 durably lowers cholesterol in primates. Nature 593, 429–434.

48. Dobosy, J.R., Rose, S.D., Beltz, K.R., Rupp, S.M., Powers, K.M., Behlke, M.A., and Walder, J.A. (2011). RNase H-dependent PCR (rhPCR): improved specificity and single nucleotide polymorphism detection using blocked cleavable primers. BMC Biotechnol. 11, 80.

49. Nahmad, A.D., Reuveni, E., Goldschmidt, E., Tenne, T., Liberman, M., Horovitz-Fried, M., Khosravi, R., Kobo, H., Reinstein, E., Madi, A., et al. (2022). Frequent aneuploidy in primary human T cells after CRISPR-Cas9 cleavage. Nat. Biotechnol. 40, 1807–1813.

50. Davis, L., Khoo, K.J., Zhang, Y., and Maizels, N. (2020). POLQ suppresses interhomolog recombination and loss of heterozygosity at targeted DNA breaks. Proc. Natl. Acad. Sci. U. S. A. 117, 22900–22909.

51. Boutin, J., Rosier, J., Cappellen, D., Prat, F., Toutain, J., Pennamen, P., Bouron, J., Rooryck, C., Merlio, J.P., Lamrissi-Garcia, I., et al. (2021). CRISPR-Cas9 globin editing can induce megabase-scale copy-neutral losses of heterozygosity in hematopoietic cells. Nat. Commun. 12, 4922.

52. Schiroli, G., Conti, A., Ferrari, S., Della Volpe, L., Jacob, A., Albano, L., Beretta, S., Calabria, A., Vavassori, V., Gasparini, P., et al. (2019). Precise Gene Editing Preserves Hematopoietic Stem Cell Function following Transient p53-Mediated DNA Damage Response. Cell Stem Cell 24, 551–565.e8.

53. Maganti, H.B., Bailey, A.J.M., Kirkham, A.M., Shorr, R., Pineault, N., and Allan, D.S. (2021). Persistence of CRISPR/Cas9 gene edited hematopoietic stem cells following transplantation: A systematic review and meta-analysis of preclinical studies. Stem Cells Transl. Med. 10, 996–1007.

54. Gothot, A., van der Loo, J.C., Clapp, D.W., and Srour, E.F. (1998). Cell cycle-related changes in repopulating capacity of human mobilized peripheral blood CD34(+) cells in non-obese diabetic/severe combined immune-deficient mice. Blood 92, 2641–2649.

55. Uchida, N., He, D., Friera, A.M., Reitsma, M., Sasaki, D., Chen, B., and Tsukamoto, A. (1997). The unexpected G0/G1 cell cycle status of mobilized hematopoietic stem cells from peripheral blood. Blood 89, 465–472.

56. Hustedt, N., and Durocher, D. (2016). The control of DNA repair by the cell cycle. Nat. Cell Biol. 19, 1–9.

57. Suzuki, K., Tsunekawa, Y., Hernandez-Benitez, R., Wu, J., Zhu, J., Kim, E.J., Hatanaka, F., Yamamoto, M., Araoka, T., Li, Z., et al. (2016). In vivo genome editing via CRISPR/Cas9 mediated homology-independent targeted integration. Nature 540, 144– 149.

58. Kim, S., Kim, D., Cho, S.W., Kim, J., and Kim, J.-S. (2014). Highly efficient RNA-guided genome editing in human cells via delivery of purified Cas9 ribonucleoproteins. Genome Res. 24, 1012–1019.

59. Kramara, J., Osia, B., and Malkova, A. (2018). Break-Induced Replication: The Where, The Why, and The How. Trends Genet. 34, 518–531.

60. Heddle, J.A., and Carrano, A.V. (1977). The DNA content of micronuclei induced in mouse bone marrow by γ-irradiation: evidence that micronuclei arise from acentric chromosomal fragments. Mutat. Res./Fundam. Mol. Mech. Mutag. 44, 63–69.

61. Liu, S., and Pellman, D. (2020). The coordination of nuclear envelope assembly and chromosome segregation in metazoans. Nucleus 11, 35–52.

62. Lauridsen, F.K.B., Jensen, T.L., Rapin, N., Aslan, D., Wilhelmson, A.S., Pundhir, S., Rehn, M., Paul, F., Giladi, A., Hasemann, M.S., et al. (2018). Differences in Cell Cycle Status Underlie Transcriptional Heterogeneity in the HSC Compartment. Cell Rep. 24, 766–780.

63. Oedekoven, C.A., Belmonte, M., Bode, D., Hamey, F.K., Shepherd, M.S., Che, J.L.C., Boyd, G., McDonald, C., Belluschi, S., Diamanti, E., et al. (2021). Hematopoietic stem cells retain functional potential and molecular identity in hibernation cultures. Stem Cell Reports 16, 1614–1628.

64. Hoban, M.D., Cost, G.J., Mendel, M.C., Romero, Z., Kaufman, M.L., Joglekar, A.V., Ho, M., Lumaquin, D., Gray, D., Lill, G.R., et al. (2015). Correction of the sickle cell disease mutation in human hematopoietic stem/progenitor cells. Blood 125, 2597–2604.

65. DeWitt, M.A., Magis, W., Bray, N.L., Wang, T., Berman, J.R., Urbinati, F., Heo, S.-J., Mitros, T., Muñoz, D.P., Boffelli, D., et al. (2016). Selection-free genome editing of the sickle mutation in human adult hematopoietic stem/progenitor cells. Sci. Transl. Med. 8, 360ra134.

66. Dever, D.P., Bak, R.O., Reinisch, A., Camarena, J., Washington, G., Nicolas, C.E., Pavel-Dinu, M., Saxena, N., Wilkens, A.B., Mantri, S., et al. (2016). CRISPR/Cas9 β- globin gene targeting in human haematopoietic stem cells. Nature 539, 384–389.

67. Shin, J.J., Schröder, M.S., Caiado, F., Wyman, S.K., Bray, N.L., Bordi, M., Dewitt, M.A., Vu, J.T., Kim, W.-T., Hockemeyer, D., et al. (2020). Controlled Cycling and Quiescence Enables Efficient HDR in Engraftment-Enriched Adult Hematopoietic Stem and Progenitor Cells. Cell Rep. 32, 108093.

68. Philippidis, A. (2023). Graphite Bio Pauses Lead Gene Editing Program in Sickle Cell Disease. Hum. Gene Ther. 34, 90–93.

69. Brendel, C., Negre, O., Rothe, M., Guda, S., Parsons, G., Harris, C., McGuinness, M., Abriss, D., Tsytsykova, A., Klatt, D., et al. (2020). Preclinical Evaluation of a Novel Lentiviral Vector Driving Lineage-Specific BCL11A Knockdown for Sickle Cell Gene Therapy. Mol Ther Methods Clin Dev 17, 589–600.

70. Yardeni, T., Eckhaus, M., Morris, H.D., Huizing, M., and Hoogstraten-Miller, S. (2011). Retro-orbital injections in mice. Lab Anim. 40, 155–160.

71. Narina, S., Connelly, J.P., and Pruett-Miller, S.M. (2023). High-Throughput Analysis of CRISPR-Cas9 Editing Outcomes in Cell and Animal Models Using CRIS.py. Methods Mol. Biol. 2631, 155–182.

72. Connelly, J.P., and Pruett-Miller, S.M. (2019). CRIS.py: A Versatile and High-throughput Analysis Program for CRISPR-based Genome Editing. Sci. Rep. 9, 4194.

73. Buenrostro, J.D., Giresi, P.G., Zaba, L.C., Chang, H.Y., and Greenleaf, W.J. (2013). Transposition of native chromatin for fast and sensitive epigenomic profiling of open chromatin, DNA-binding proteins and nucleosome position. Nat. Methods 10, 1213– 1218.

74. Hitz, B.C., Lee, J.-W., Jolanki, O., Kagda, M.S., Graham, K., Sud, P., Gabdank, I., Seth Strattan, J., Sloan, C.A., Dreszer, T., et al. (2023). The ENCODE Uniform Analysis Pipelines. bioRxiv. 10.1101/2023.04.04.535623.

75. Bolger, A.M., Lohse, M., and Usadel, B. (2014). Trimmomatic: a flexible trimmer for Illumina sequence data. Bioinformatics 30, 2114–2120.

76. Clement, K., Rees, H., Canver, M.C., Gehrke, J.M., Farouni, R., Hsu, J.Y., Cole, M.A., Liu, D.R., Joung, J.K., Bauer, D.E., et al. (2019). CRISPResso2 provides accurate and rapid genome editing sequence analysis. Nat. Biotechnol. 37, 224–226.

77. Liang, S.-Q., Liu, P., Smith, J.L., Mintzer, E., Maitland, S., Dong, X., Yang, Q., Lee, J., Haynes, C.M., Zhu, L.J., et al. (2022). Genome-wide detection of CRISPR editing in vivo using GUIDE-tag. Nat. Commun. 13, 437.

78. Rodríguez, T.C., Dadafarin, S., Pratt, H.E., Liu, P., Amrani, N., and Zhu, L.J. (2021). Genome-wide detection and analysis of CRISPR-Cas off-targets. Prog. Mol. Biol. Transl. Sci. 181, 31–43.

79. Zhu, L.J., Lawrence, M., Gupta, A., Pagès, H., Kucukural, A., Garber, M., and Wolfe, S.A. (2017). GUIDEseq: A bioconductor package to analyze GUIDE-Seq datasets for CRISPR-Cas nucleases. BMC Genomics 18, 1–10.

80. Tsai, S.Q., Zheng, Z., Nguyen, N.T., Liebers, M., Topkar, V.V., Thapar, V., Wyvekens, N., Khayter, C., Iafrate, A.J., Le, L.P., et al. (2015). GUIDE-seq enables genome-wide profiling of off-target cleavage by CRISPR-Cas nucleases. Nat. Biotechnol. 33, 187–197.

81. Park, J., Bae, S., and Kim, J.-S. (2015). Cas-Designer: a web-based tool for choice of CRISPR-Cas9 target sites. Bioinformatics 31, 4014–4016.

82. Doench, J.G., Fusi, N., Sullender, M., Hegde, M., Vaimberg, E.W., Donovan, K.F., Smith, I., Tothova, Z., Wilen, C., Orchard, R., et al. (2016). Optimized sgRNA design to maximize activity and minimize off-target effects of CRISPR-Cas9. Nat. Biotechnol. 34, 184–191.

83. Lowy-Gallego, E., Fairley, S., Zheng-Bradley, X., Ruffier, M., Clarke, L., Flicek, P., and 1000 Genomes Project Consortium (2019). Variant calling on the GRCh38 assembly with the data from phase three of the 1000 Genomes Project. Wellcome Open Res 4, 50.

84. Bao, X.R., Pan, Y., Lee, C.M., Davis, T.H., and Bao, G. (2021). Tools for experimental and computational analyses of off-target editing by programmable nucleases. Nat. Protoc. 16, 10–26.

85. Chaudhari, H.G., Penterman, J., Whitton, H.J., Spencer, S.J., Flanagan, N., Lei Zhang, M.C., Huang, E., Khedkar, A.S., Toomey, J.M., Shearer, C.A., et al. (2020). Evaluation of Homology-Independent CRISPR-Cas9 Off-Target Assessment Methods. CRISPR J 3, 440–453.

86. Firth, D. (1993). Bias reduction of maximum likelihood estimates. Biometrika 80, 27–38.

87. Kosmidis, I. (2017). brglm2: Bias reduction in generalized linear models. R package version 0. 1 *5*.

88. Benjamini, Y., and Hochberg, Y. (1995). Controlling the false discovery rate: A practical and powerful approach to multiple testing. J. R. Stat. Soc. 57, 289–300.

